# Nanoscale Lattice Organization of Molecular Condensates Drives Compositional Degeneracy in Synaptic Plasticity

**DOI:** 10.1101/2025.03.14.641702

**Authors:** Pallavi Rao Netrakanti, Yukti Chopra, Richa Agarwal, Premchand Rajeev, Shekhar Kedia, Mini Jose, Suhita Nadkarni, Deepak Nair

**Affiliations:** Centre for Neuroscience, Indian Institute of Science, Bangalore 560012 (India); Indian Institute of Science Education and Research, Pune 411008 (India)

## Abstract

Synaptic plasticity is essential for neuronal communication, involving coordinated structural, molecular and functional changes that are shaped by the nanoscale alterations of the active zone and postsynaptic density. Emerging evidence suggests that synapses function as complex information processing machines, where unique molecular assemblies shape transmission properties. Central to this is the organization of voltage-gated calcium channels (VGCCs) and Bassoon within active zones. Utilizing advanced techniques like liquid-liquid phase separation, super-resolution microscopy, and data-driven models of synaptic transmission, we reveal how nanoscale "compositional degeneracy" in Bassoon and VGCCs enables synapses to achieve functional adaptability through multiple molecular configurations. By modulation of local uncertainty and implementing probabilistic inference, synapses fine-tune transmission efficiency by regulating dynamic entropy and free energy. These principles are especially evident during homeostatic scaling, where synaptic scaling mechanisms differ with neuronal maturity. This study highlights how distinct thermodynamic states in VGCC and Bassoon organization optimize information transfer at different plasticity stages. Our findings propose a refined framework for understanding synaptic transmission as an adaptable, entropy-modulated process, balancing resilience and efficiency.

## Introduction

Plasticity, in its various manifestations, initiates numerous transformations in both the structural and functional aspects of synapses^1-4^. These changes encompass elements of synaptic heterogeneity, including the instantaneous availability of the correct number of molecules, permutations and combinations of signaling and conformational states of molecules, dimensions of synaptic compartments, and rates of molecular transport^2,5-8^. However, understanding these intricate alterations requires more than isolated morpho-functional analysis. We propose that each synapse incorporates a distinctive molecular framework into its lattice organization. This implies that synaptic transmission rates can be controlled by different combinations of lateral placements of synaptic molecules in specific permutations, which influence local release probability and are governed by the instantaneous state of the synapse. This lattice organization governs "nanoscale compositional degeneracy," finely tuned to optimize information transmission^9-11^. Our investigation is aimed to determine whether variations in homeostatic scaling between younger and more mature neurons are rooted in nanoscale heterogeneity within synaptic organization^12-14^. To understand the basis of molecular degeneracy, we relied on evaluating the organization of Voltage Gated Calcium Channel (VGCC) within synaptic subcompartments ^5,15-19^, we employed three phases of a pivotal analytical framework that examines synapse as an information processing machine. Each phase is built upon the insights of the previous one, addressing whether individual synapses function as information transfer domains as indicated below:

1. **Representation of uncertainty and its impact on information processing**: In this initial stage, we investigated the nanoscale segregation of VGCCs—a functional marker of homeostatic scaling at the presynapse—and Bassoon—a structural marker for the cytomatrix of the active zone. Applying principles of liquid-liquid phase separation in an N-component system, we modeled synaptic organization as a first-order phase transition and extracted relative changes in entropy between hierarchical organization of molecules^20,21^.
2. **Neural implementation of probabilistic inference and its role in homeostatic plasticity**: Building on the previous phase, we explored whether VGCCs, due to their direct influence on intracellular calcium levels, exhibit distinctive signatures of synaptic scaling. We assessed signatures of multiplicative scaling that remained consistent across diverse functional zones of excitatory synapses^22,23^.
3. **Spatial organization of molecular condenstates and and its Impact on synaptic scaling**: In the final phase, we conducted an in-depth evaluation of organizational variability within presynaptic compartments. Using multi-parameter image analysis, we extracted 14 morpho-functional parameters defining molecular organization from a single region of interest (28 from a single synapse), generating a Cartesian geometry of lattice organization for independent nanodomains of VGCCs, Bassoon, and their co-condensates. This comprehensive approach allowed us to analyze hundreds of synapses and active zones using multi-parameter analysis of super-resolved images. The observed structural heterogeneity within individual excitatory presynaptic compartments was used in data-driven models of synaptic transmission to understand how nanoscale degeneracy in molecular organization impacts synaptic transmission.

With the above-mentioned framework, we present compelling evidence for modeling chemical synaptic transmission within an information processing regime that integrates the representation of uncertainty, its implementation in neural code, and spatial basis functions^24-26^. This approach helps us to elucidate the differences between structural rules governing various rates of synaptic transmission at the nanoscale, where subtle changes in modular assembly can be a powerful tool for altering synaptic properties.

## Results

### Neuronal development alters thermodynamic fingerprints of nanoscale condensates and differentially regulate structural and functional markers of synaptic transmission

Although spontaneous and action potential evoked release differ in their spatial and functional properties, evidence supports the involvement of a common subset of molecules in both of these events^6,27,28^. This is evident in paradigms that evoke synaptic scaling, a form of homeostatic plasticity that is induced by the absence of action potential-dependent release events^13,14^. A recent study found that P/Q-type VGCC (also known as CaV2.1) is required for presynaptic homeostatic plasticity, regulating basal neurotransmission and synaptic vesicle pool size^15-17^. As localization and molecular state of VGCC directly influences local calcium concentration to alter release probability, we denoted CaV2.1 α-subunit of the P/Q-type VGCC as a functional marker^17,29^. Bassoon, a cardinal protein of the cytomatrix necessary for the integrity of the cytomatrix of the active zone and for VGCC localization, is denoted as structural marker^16,30^ (Fig. S1, Scheme 1). To gain a better understanding of the relationship between nanoscale structure and function, we initiated an investigation into the heterogeneity of chemical information processing arising from the liquid-liquid phase separation of molecules in the cytomatrix of the active zone (AZ). In analyzing the phase transition in an N-component system such as synapses, it is necessary to correlate sequential variability in copy numbers of proteins in spatially segregated functional zones of synapses^21,31^. To evaluate molecular parameters describing the heterogeneous distribution of these molecules within the functional zones of the synapse, we employed both single molecule and ensemble based super-resolution techniques (Fig. 1A, D, G, J and Fig. S1, Fig. S2, S3). Immunolabelled primary hippocampal neurons for Bassoon and VGCC were first imaged at the diffraction limit using Highly Inclined and Laminated Optical sheet (HILO) illumination and then using Direct Stochastic Optical Reconstruction Microscopy (dSTORM). Examination of synapses and active zones using HILO illumination revealed a diffused distribution of Bassoon and VGCC at both DIV 8 and DIV 15 (Fig. 1A-i, Fig. 1G-i and Fig. 1D-i, Fig. 1J-i). The diffraction-limited image of the Bassoon enriched compartment served as a presynaptic marker, while the super-resolution image of the same was used as a marker for the AZ. Although the majority of Bassoon in the synaptic compartment was enriched in the AZ, the distribution of Bassoon in the AZ itself displayed high variability and increased packing density. Super-resolution imaging by dSTORM showed that both Bassoon and VGCC exhibit punctate distribution within the AZ and the presynaptic compartment (Fig. 1A-ii, 1G-ii for Bassoon, and Fig. 1D-ii, Fig. 1J-ii for VGCC at DIV 8 and DIV 15, respectively). We then evaluated the heterogeneity in the distribution of structural and functional markers in synapses of neurons of DIV 8 and 15 upon induction of homeostatic plasticity. DIV 8 was chosen because it coincides with the peak of synaptogenesis, while DIV 15 correlated with mature stages of synapses, displaying postsynaptic pruning^12^. We refer to 24 hours post TTX application at DIV 8 as Early Homeostatic Scaling (EHS) and the 24 hours post TTX application at DIV 15 as Late Homeostatic Scaling (LHS). The same cultures were used longitudinally, while untreated neurons from the same biological replicate at DIV 8 and DIV 15 were used as corresponding controls.

**Figure 1:**
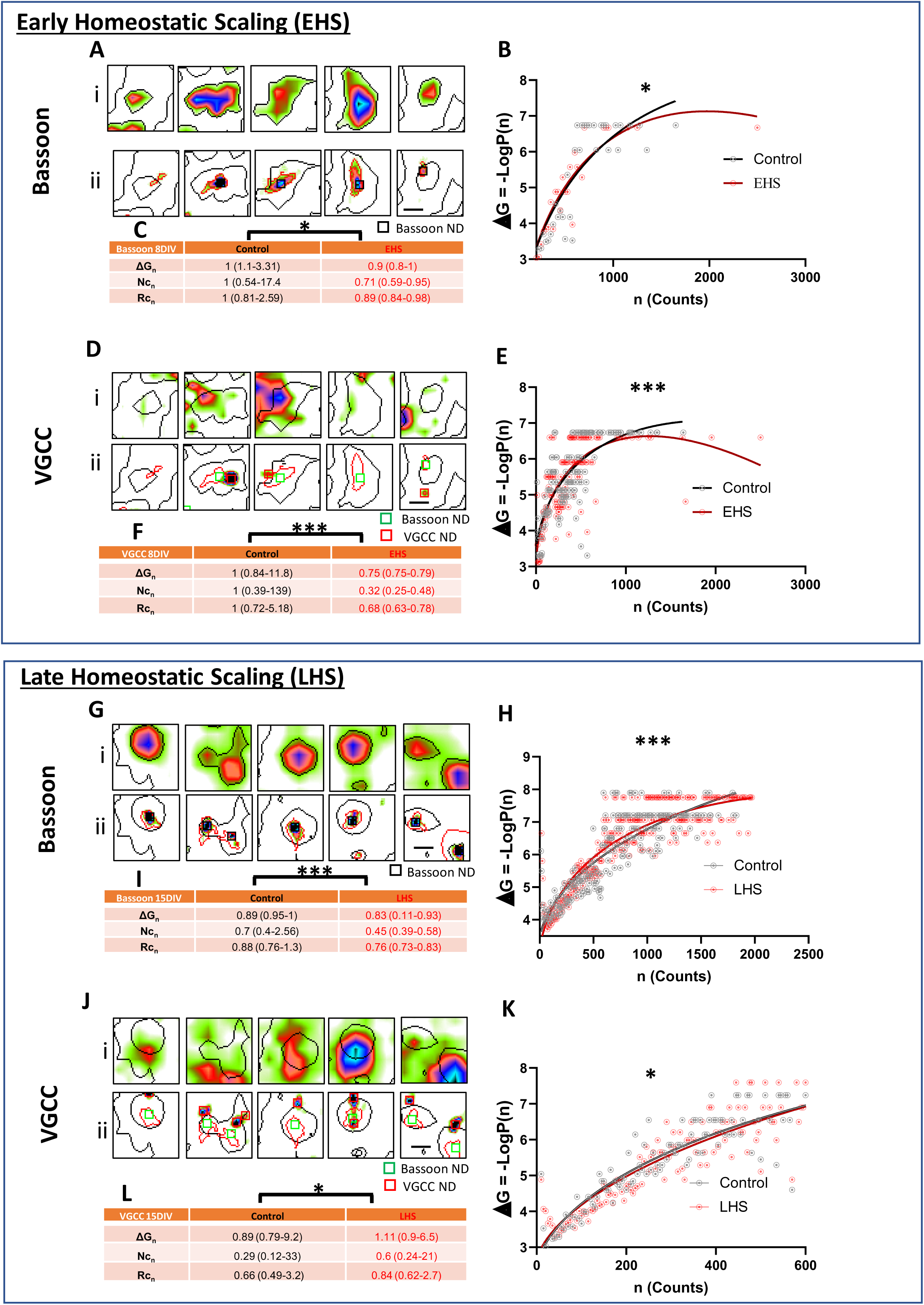
Nanoscale organization of Bassoon and VGCC follows a first-order phase transition in early and late homeostatic scaling: (A) (i) Pseudocolor images of Bassoon distribution in synapses at DIV 8. Bassoon regions were automatically detected by thresholding the image and are marked by black contours, representing synaptic regions. (ii) Pseudocolor images of Bassoon nanodomains (black squares) within synapses. (B) Free energy curve fit of the probability distribution of Bassoon molecules at DIV 8 for control and EHS. (C) Table summarizing ΔG, Nc, and Rc of Bassoon at DIV 8 between control and EHS. (D) (i) Pseudocolor images of VGCC distribution in synapses at DIV 8. (ii) Pseudocolor images of VGCC nanodomains (red squares) and Bassoon (green squares) within synapses. (E) Free energy curve fit of the probability distribution of VGCC molecules at DIV 8 for control and EHS. (F) Table summarizing ΔG, Nc, and Rc of VGCC at DIV 8 between control and EHS. All the values are normalised to EHS control (G) (i) Pseudocolor images of Bassoon distribution in synapses at DIV 15. (ii) Pseudocolor images of Bassoon nanodomains (black squares) within synapses. (H) Free energy curve fit of the probability distribution of Bassoon molecules at DIV 15 for control and LHS. (I) Table summarizing the ΔG, Nc, and Rc of Bassoon at DIV 15 between control and LHS. All the values are normalised to EHS control (J) (i) Pseudocolor images of VGCC distribution in synapses at DIV 15. (ii) Pseudocolor images of VGCC nanodomains (red squares) and Bassoon nanodomains (green squares) within synapses. (K) Free energy curve fit of the probability distribution of VGCC molecules at DIV 15 for Control and LHS. (L) Table summarizing the ΔG, Nc, and Rc of VGCC at DIV 15 between control and LHS. All the values are normalised to EHS control. The n values for the free energy curve fit, ΔG, Nc, and Rc of Bassoon at DIV 8 are control: 849, EHS: 743; and at DIV 15, control: 2695, LHS: 2333. The n values for the free energy curve fit, ΔG, Nc, and Rc of VGCC at DIV 8 are control: 847, EHS: 738; and at DIV 15, control: 1400, LHS: 1992.

To extract the molecular traits of single order phase transitions resulting from the induction of homeostatic plasticity via synaptic scaling, we relied on classical nucleation theory and the Szilard model of non-equilibrium steady state supersaturation ^21,31^. Using dSTORM microscopy, we probed the parameters associated with single order phase transitions for Bassoon and VGCC in DIV 8 and DIV 15 neurons (Fig. S4 A-E). We extracted the copy number of detected Bassoon/VGCC per nanodomain, which were observed as discrete puncta in the synaptic compartment. The copy number thus extracted from synaptic compartments served as the raw data for phase transition analysis (see Materials and Methods). The free energy change in assembling a cluster of n molecules from the disperse phase is given by the equation ΔG = an2/3 ± bn + c. This equation comprises of two major elements: surface energy and bulk energy. In phase separation, the interface between two liquid phases of demixed components represents the surface energy (ΔG surface = an2/3). The difference in free energy between a system with all molecules in the ambient phase and a system with n molecules in the clustered phase represents the bulk energy (bn).

The slope of the bulk energy (bn) was negative for Bassoon and VGCC during both EHS and LHS (Fig. S5), which is indicative of a supersaturated system. We performed a free energy curve fit of the probability distribution of Bassoon (Fig. 1B) and VGCC molecules (Fig. 1 E) in control versus EHS (TTX), allowing us to obtain the parameters a, b, and c. These parameters helped us to calculate the nucleation barrier (Nc) and critical cluster radius (Rc). Tabulated values of ΔG, Nc, and Rc for Bassoon and VGCC are shown in Fig. 1 C and 1 F, respectively. During EHS, Bassoon and VGCC tend to cluster more readily, as indicated by the reduced ΔG, Nc, and Rc values for Bassoon and the negative slope of bn for both proteins (Fig. S5 A-D). Similarly, VGCC tend to cluster more easily than control during EHS (Fig. 1 F). Interestingly, at DIV 15, the free energy curves for both Bassoon and VGCC, along with their ΔG, Nc, and Rc values were significantly different between control and TTX treated cells. This indicates that homeostatic scaling regulates the phenomenon of cluster formation differently for Bassoon and VGCC depending on the age of the neuron. Moreover, the negative slopes for bn for both proteins were similar between LHS and EHS, indicating a supersaturated system at DIV 15 (Fig. S5 E-H).

### Multiplicative scaling of structural and functional markers during homeostatic plasticity is conserved but non-uniform within the presynaptic compartment

We initially analyzed the thermodynamic variables of the first-order phase transition for Bassoon and VGCC, and we elucidated the molecular signatures for their supersaturated state leading to clusters (Fig. 1). The next step of investigation was aimed to determine whether the nanoscale condensates of Bassoon and VGCC were altered and undergo a multiplicative change during TTX induced homeostatic plasticity paradigm. To achieve this, we focused our efforts on understanding the spatial organization and quantitative levels of Bassoon and VGCC in the AZ and in different functional compartments of synapses.To accomplish this, we utilized the ensemble nature of super-resolution imaging by Stimulated Emission Depletion (STED) Microscopy, which allowed us to characterize Bassoon and VGCC in synapses and AZ in detail. We also employed sequential confocal and STED microscopy to evaluate the diffraction-limited and nanoscale distribution of these proteins. Comparable paradigms of imaging have been successfully used to evaluate the contribution of associated synaptic molecules that localize to different functional zones of the synapse^23,32,33^.

The distributions of Bassoon and VGCC in neuronal processes, synapses, and AZ were examined using confocal and STED microscopy and were found comparable to observations by HILO and dSTORM (Fig. 2 A-D, Fig. 2E-H, Fig. 3 A-D, Fig. 3 E-H). We focused on VGCC for the initial part of this analysis. During EHS, an increase in the overall levels of VGCC was observed in terms of average intensity at the synapse, AZ, and nanodomains within the AZ (Fig. 2 I, L, and R), while nanodomains in the whole presynaptic compartment remained unaffected (Fig. 2 O). As VGCC is a functional marker and directly responsible for the synaptic concentration of Ca^2+^, the copy number of VGCC was investigated to determine if it displayed multiplicative changes similar to homeostatic scaling^23^. A rank-order analysis was conducted on the average intensity values of both control and TTX datasets (Fig. S6 A-D). For this, the intensities were sorted in ascending order, and a linear fit was calculated after plotting the TTX dataset against the control distribution. The slope of the linear fit was used to estimate the scaling factor (Fig. S6 A-D). To determine the accurate scaling factor, the TTX dataset was scaled with various arbitrary scaling factors, and the scaled datasets were compared to the control dataset using the Kolmogorov–Smirnov (K-S) test. The scaling factor corresponding to the highest p-value was considered the most accurate. The data revealed that average intensity of VGCC in the synapse and AZ and in nanodomains of AZ during EHS exhibited multiplicative scaling (Fig. 2 J, M, and S), while the nanodomains of VGCC in the whole presynaptic compartment remained similar to that of the control (Fig. 2 P). Upon plotting the cumulative frequency distribution of the scaled data (obtained by dividing the EHS (TTX) data with its corresponding scaling value), along with EHS and control data, it was evident that the scaled TTX distribution overlapped with the control distribution (Fig. 2 K, N). This demonstrated successful multiplicative scaling of VGCC numbers in both synapse and AZ when homeostatic scaling was activated in EHS. However, multiplicative scaling of VGCC nanodomains was not observed in the synaptic compartment (Fig. 2 Q). Nevertheless, a significant change was observed for VGCC nanodomains in the AZ (Fig. 2 T). A similar analysis of VGCC during LHS confirmed its increased content in synapses and AZ. Moreover, there was an increase in VGCC content in nanodomains in both synapses and AZ (Fig. 3I, L, O, and R). During LHS, multiplicative scaling of VGCC was observed in both synapses (Fig. 3J, K) and AZ (Fig. 3 M, N) and in nanodomains of these compartments (synapses - Fig. 3 P, Q, AZ - Fig. 3 S, T). Additionally, the multiplicative nature of VGCC in discrete spatial zones of the synapse was confirmed by a simple linear fit of the rank-ordered data during EHS and LHS (Fig. S7 A-D, Supp Table 1). These observations confirm that the multiplicative scaling of VGCC is conserved but not identical across subsynaptic compartments and nanodomains.

**Figure 2:**
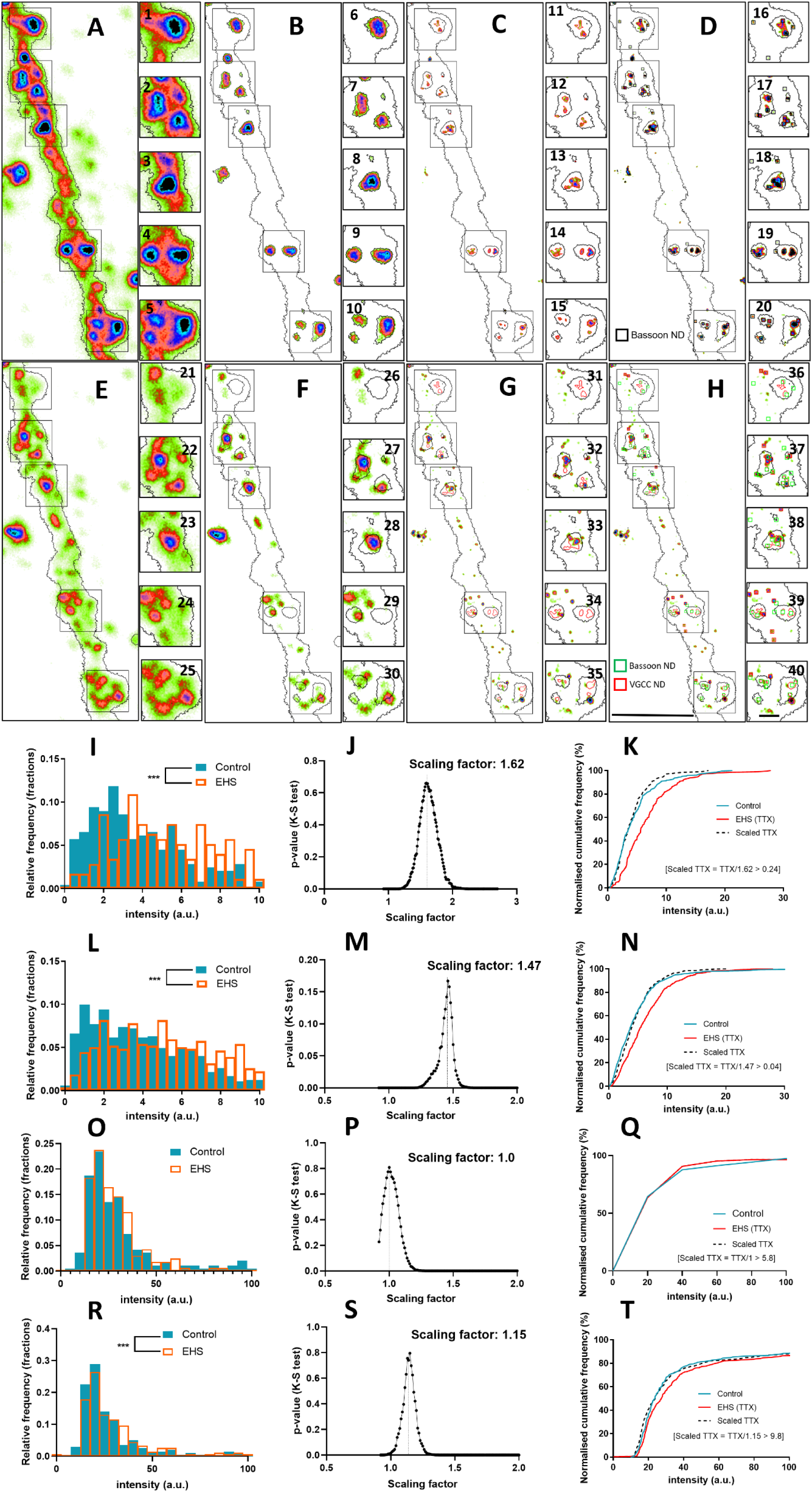
Effect of early homeostatic scaling on VGCC average intensity in synapses and active zones (AZ): (A) Pseudocolor epifluorescence image of Bassoon in a DIV 8 neuron, with black contours indicating automatically detected neuronal processes. Insets 1–5 show potential synapses. (B) Pseudocolor epifluorescence image of Bassoon in the neuron, with black contours (automatically detected confocal Bassoon regions) indicating synapses. Insets 6–10 show Bassoon distribution within synapses. (C) Pseudocolor STED image of Bassoon in the same neuron, with red contours within the black Bassoon confocal contours indicating automatically detected STED regions, representing AZs. Insets 11–15 show Bassoon distribution within AZs. (D) STED image of Bassoon in the same neuron, with black squares indicating automatically detected Bassoon clusters. Insets 16–20 show detected Bassoon nanodomains. (E) Pseudocolor epifluorescence image of VGCC in a DIV 8 neuron, with black contours indicating neuronal processes. Insets 21–25 show potential synapses. (F) Pseudocolor epifluorescence image of VGCC in the neuron, with black contours (automatically detected confocal VGCC regions) marking synapses. Insets 26–30 show VGCC distribution within synapses. (G) Pseudocolor STED image of VGCC in the same neuron. Insets 31–35 show VGCC distribution within AZs. (H) Pseudocolor STED image of VGCC in the same neuron, with green squares indicating Bassoon clusters and red squares indicating VGCC clusters. Insets 36–40 show nanodomains of Bassoon and VGCC. Scale bar in (H) represents 1 µm, and insets correspond to 0.5 µm. ND = Nanodomain. (I) VGCC average intensity per synapse in a DIV 8 neuron under control and EHS conditions (n=267 synapses for control, n=210 for TTX dataset). (J) The EHS dataset was scaled and compared with control, with a scaling factor for VGCC in synapses at DIV 8 of 1.62 (p=0.658, K-S test). (K) Normalized cumulative frequency distribution of control, EHS (TTX), and scaled TTX for VGCC intensity in synapses at DIV 8, showing no significant difference between scaled TTX and control data.(L) VGCC average intensity per AZ at DIV 8 between control and EHS conditions (n=703 AZ for control, n=631 for TTX dataset). (M) Scaling factor for VGCC in AZs at DIV 8 (1.47, p=0.167, K-S test). (N) Normalized cumulative frequency distribution of control, EHS (TTX), and scaled TTX for VGCC intensity in AZs at DIV 8, showing no significant difference between scaled TTX and control data.(O) VGCC nanodomain average intensity per synapse in DIV 8 neurons under control and EHS conditions (n=197 synapses for control, n=170 for EHS dataset). (P) Scaling factor for VGCC nanodomains in synapses at DIV 8 (1.0, p=0.809, K-S test). ( Q) Normalized cumulative frequency distribution of control, EHS (TTX), and scaled TTX for VGCC nanodomain intensity in synapses at DIV 8. (R) VGCC nanodomain average intensity per AZ in DIV 8 neurons under control and EHS conditions (n=226 AZ for control, n=309 for EHS dataset). (S) Scaling factor for VGCC nanodomains in AZs at DIV 8 (1.15, p=0.795, K-S test). (T) Normalized cumulative frequency distribution of control, EHS (TTX), and scaled TTX for VGCC nanodomain intensity in AZs at DIV 8.

**Figure 3:**
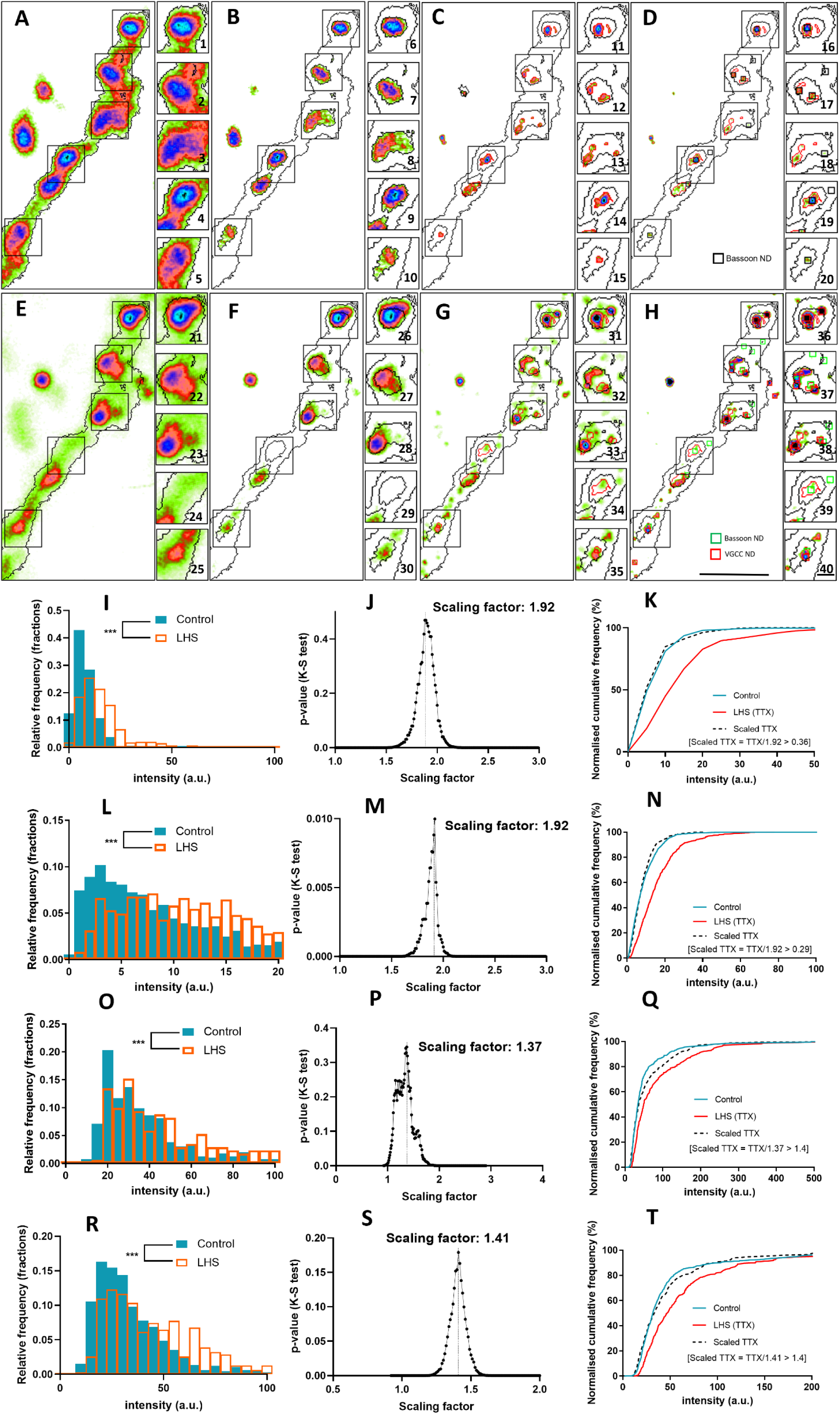
Effect of late homeostatic scaling on the average intensity of VGCC in synapses and active zones (AZ). (A) Pseudocolor epifluorescence image of Bassoon in a DIV 15 neuron, where the black contour indicates the automatic detection of neuronal processes. Insets 1-5 display a gallery of potential synapses. (B) Pseudocolor epifluorescence image of Bassoon in the same DIV 15 neuron, where the black contour (automatically detected confocal Bassoon regions) indicates synapses. Insets 6-10 show the distribution of Bassoon within these synapses. (C) Pseudocolor STED image of Bassoon in the same neuron, where the red contour within the black Bassoon confocal contour represents automatically detected STED regions of Bassoon, indicating the active zone (AZ) region. Insets 11-15 depict the distribution of Bassoon within the AZ. (D) Pseudocolor STED image of Bassoon in the same neuron, where the black squares represent automatically detected clusters of Bassoon. Insets 16-20 illustrate the detected nanodomains of Bassoon. (E) Pseudocolor epifluorescence image of VGCC in the DIV 15 neuron, where the black contour indicates the automatic detection of neuronal processes. Insets 21-25 show a gallery of potential synapses. (F) Pseudocolor epifluorescence image of VGCC in the same neuron, where the black contour (automatically detected confocal Bassoon regions) indicates synapses. Insets 26-30 present the distribution of VGCC within these synapses. (G) Pseudocolor STED image of VGCC in the same neuron. Insets 31-35 illustrate the distribution of VGCC within the AZ. (H) Pseudocolor STED image of VGCC in the same neuron, where the green squares represent automatically detected clusters of Bassoon, and the red squares represent automatically detected clusters of VGCC. Insets 36-40 depict the nanodomains of both Bassoon and VGCC. The scale bars in H represent 1 micron, and the insets correspond to 0.5 microns. ND: Nanodomain. (I) Distribution of VGCC average intensity per synapse in DIV 15 neurons under control and LHS conditions (n=627 and 372 synapses for control and LHS datasets). (J) Multiplicative scaling factor for VGCC in the synapse at DIV 15 (1.92, p=0.448, K-S test). (K) Normalized cumulative frequency distribution of control, LHS (TTX), and scaled TTX for VGCC average intensity in synapses at DIV 15. The scaled TTX distribution was not significantly different from the control. (L) Distribution of VGCC average intensity per AZ in DIV 15 neurons under control and LHS conditions (n=1386 and 825 AZ for control and LHS datasets). (M) Multiplicative scaling factor for VGCC in the AZ at DIV 15 (1.92, p=0.00998, K-S test). (N) Normalized cumulative frequency distribution of control, LHS (TTX), and scaled TTX for VGCC average intensity in AZ at DIV 15. The scaled TTX distribution was not significantly different from the control. (O) Distribution of the average intensity of VGCC nanodomains per synapse in DIV 15 neurons under control and LHS conditions (n=449 and 227 synapses for control and LHS datasets). (P) The LHS dataset was scaled using multiple values and compared with the control. The scaling factor providing the maximum p-value between the scaled TTX and control datasets was chosen as the multiplicative scaling factor for average intensity of VGCC nanodomains in the synapse at DIV 15 (1.37, p=0.345, K-S test).(Q) Normalized cumulative frequency distribution of control, LHS (TTX), and scaled TTX for average intensity of VGCC nanodomains in synapses at DIV 15. (R) Distribution of average intensity of VGCC nanodomains per AZ in DIV 15 synapses under control and LHS conditions (n=1386 and 825 AZ for control and LHS datasets). (S) The LHS dataset was scaled using multiple values and compared with the control. The scaling factor providing the maximum p-value between the scaled TTX and control datasets was chosen as the multiplicative scaling factor for average intensity of VGCC nanodomains in the AZ at DIV 15 (1.41, p=0.176, K-S test).(T) Normalized cumulative frequency distribution of control, LHS (TTX), and scaled TTX for average intensity of VGCC nanodomains in AZ at DIV 15.

Spatially conserved, non-identical signatures of regulation exist within subsynaptic zones that are modulated by a differential assembly of molecules. We evaluated if such changes are associated with molecules that serve as structural scaffolds to functional markers like VGCC. To validate if such changes were directly correlated with Bassoon, we evaluated the propensity of Bassoon to undergo multiplicative scaling. The molecular content of Bassoon increased in both the synapse and AZ during EHS and LHS (Fig. S8 A, D, and G, J). Although there was a significant increase in the molecular content of Bassoon, it did not follow the pattern of multiplicative scaling in these compartments for both EHS and LHS (Fig. S8 B, C, E, F, and H, I, K, L). Similarly, even though the amount of Bassoon increased within its nanodomains in synapses and AZ during EHS and LHS (Fig. S9 A, D, and G, J), this increase was not reflective of signatures of multiplicative scaling (Supp Fig. 9 B, C, E, F, and H, I, K, L).

Since the thermodynamic changes were assessed by single-molecule localization microscopy and multiplicative scaling performed using ensemble super-resolution microscopy, we analysed if using the different techniques resulted in comparable outcomes. We extracted and compared the number of molecules in the nanodomains of the AZ using intensity ratios from the dSTORM and STED data. At DIV 15, we observed an increase in the number of molecules in nanodomains of the synapse and AZ for both Bassoon (Fig. S10 A, C) and VGCC (Fig. S10 B, D) in the dSTORM data. This increase in the number of molecules in nanodomains was consistent with the increase in the content of molecules within nanodomains in the synapses and AZ for both Bassoon (Fig. S10 E, G) and VGCC (Fig. S10 F, H) in the STED data. Our findings confirmed that the changes induced by homeostatic synaptic scaling for Bassoon and VGCC in synapses and the AZ is robust as reflected by comparable outcomes from these diverse super-resolution imaging techniques.

### Heterogeneity in cartesian geometry of lattice organization and co-condensation of structural and functional markers between EHS and LHS

As a third step and for proof of concept, we established reference frames by employing nanoscale spatial basis functions to construct a Cartesian graph of nanodomains that adhered to the principles of first-order phase transitions. We then applied multivariate analysis to comprehend the corresponding alterations in nanoscale organization, creating a structural framework for interpreting the variations in homeostatic scaling during different developmental stages. Towards this end, we identified a set of direct and indirect parameters from every synapse that corresponded to integrated morphometry analysis of structural and functional markers, enabling a multiparameter comparison to evaluate the alterations in the synaptic organization in EHS and LHS. We generated a scheme to extract 14 classifiers (Table 1-4) for any molecule of interest from a region representing the synapse or diverse functional zones of the synapse (Fig. S11).

**Table 1:** Data summarising the distribution of Bassoon in synapses during EHS and LHS.

|  | Early Homeostatic Scaling |  | Late Homeostatic Scaling |  |
| --- | --- | --- | --- | --- |
| Synaptic bouton | Bassoon Control (n=1175) | Bassoon EHS (n=833) | Bassoon Control (n=2567) | Bassoon LHS (n=2224) |
| Synaptic region average intensity (Bassoon confocal mask) (normalised to control) | $1 \pm 0.028$ (1, 0.6726 - 2.526) | $1.366 \pm 0.030$ (2.1, 1.550 - 2.931) | $1 \pm 0.015$ (1, 0.501 - 1.303) | $1.463 \pm 0.021$ (1.171, 0.773 - 1.836) |
| Synaptic region total intensity (Bassoon confocal mask) (normalised to control) | $1 \pm 0.058$ (1, 0.3965 - 2.974) | $1.486 \pm 0.076$ (2.128, 0.9166 - 6.004) | $1 \pm 0.038$ (1, 0.167 - 0.973) | $1.626 \pm 0.053$ (0.751, 0.263 - 1.866) |
| Synaptic area (Bassoon confocal mask) | $0.068 \pm 0.002$ (0.043, 0.016 - 0.089) | $0.078 \pm 0.003$ (0.050, 0.020 - 0.112) | $0.086 \pm 0.002$ (0.058, 0.024 - 0.121) | $0.099 \pm 0.002$ (0.072, 0.029 - 0.146) |
|  | Nanodomain (n = 1347) | Nanodomain (n = 898) | Nanodomain (n = 2830) | Nanodomain (n = 2532) |
| Principal axis | $0.087 \pm 0.001$ (0.082, 0.070 - 0.097) | $0.120 \pm 0.002$ (0.100, 0.085 - 0.128) | $0.140 \pm 0.002$ (0.120, 0.094 - 0.162) | $0.160 \pm 0.004$ (0.140, 0.100 - 0.180) |
| Average intensity (normalised to control) | $1 \pm 0.026$ (1, 0.6743 - 2.058) | $1.204 \pm 0.030$ (1.582, 1.020 - 2.375) | $1 \pm 0.040$ (1, 0.4505 - 1.904) | $1.325 \pm 0.019$ (1.590, 0.9682 - 2.403) |
| Total intensity (normalised to control) | $1 \pm 0.029$ (1, 0.5734 - 2.244) | $1.337 \pm 0.033$ (1.773, 1.194 - 2.812) | $1 \pm 0.017$ (1, 0.5595 - 1.886) | $1.356 \pm 0.018$ (1.532, 1.012 - 2.407) |
| Area | $0.0053 \pm 0.0001$ (0.0047, 0.0034 - 0.0064) | $0.0087 \pm 0.0002$ (0.0070, 0.0050 - 0.0104) | $0.0120 \pm 0.0003$ (0.0097, 0.0056 - 0.0160) | $0.0150 \pm 0.0006$ (0.0112, 0.0064 - 0.0180) |
| Individual intensity ratio ($IR_i$) | $0.350 \pm 0.009$ (0.268, 0.136 - 0.511) | $0.308 \pm 0.008$ (0.240, 0.134 - 0.436) | $0.222 \pm 0.006$ (0.157, 0.102 - 0.264) | $0.211 \pm 0.008$ (0.131, 0.091 - 0.234) |
| Grouped intensity ratio ($IR_g$) | $0.590 \pm 0.008$ (0.573, 0.451 - 0.698) | $0.567 \pm 0.008$ (0.542, 0.407 - 0.676) | $0.430 \pm 0.006$ (0.400, 0.282 - 0.558) | $0.446 \pm 0.008$ (0.451, 0.288 - 0.589) |
| Area ratio | $0.083 \pm 0.003$ (0.054, 0.029 - 0.106) | $0.112 \pm 0.004$ (0.076, 0.041 - 0.137) | $0.217 \pm 0.005$ (0.141, 0.059 - 0.334) | $0.256 \pm 0.008$ (0.143, 0.059 - 0.365) |
| Inter cluster distance ($Dist_{IC}$) | $0.265 \pm 0.005$ (0.256, 0.159 - 0.369) | $0.230 \pm 0.005$ (0.211, 0.133 - 0.323) | $0.220 \pm 0.004$ (0.202, 0.116 - 0.312) | $0.205 \pm 0.005$ (0.187, 0.104 - 0.284) |
| Distance from ROI centre ($Dist_{ROIc}$) | $0.087 \pm 0.002$ (0.067, 0.034 - 0.129) | $0.089 \pm 0.002$ (0.074, 0.036 - 0.134) | $0.119 \pm 0.003$ (0.096, 0.055 - 0.172) | $0.117 \pm 0.003$ (0.100, 0.050 - 0.167) |
| Distance between Bassoon and VGCC nanodomains ($Dist_{BV}$) | $0.08037 \pm 0.00397$ (0.08191, 0.05219 - 0.1081) | $0.08826 \pm 0.003793$ (0.09642, 0.0496 - 0.1246) | $0.07792 \pm 0.002244$ (0.07524, 0.04146 - 0.1155) | $0.07777 \pm 0.003754$ (0.07511, 0.04435 - 0.1098) |
| Fraction of ROIs with cluster numbers 1 to 10 | $1.674 \pm 0.04223$ (1, 1 - 2) | $1.717 \pm 0.04991$ (1, 1 - 2) | $1.907 \pm 0.03429$ (1, 1 - 2) | $2.011 \pm 0.03799$ (2, 1 - 3) |

**Table 2:**
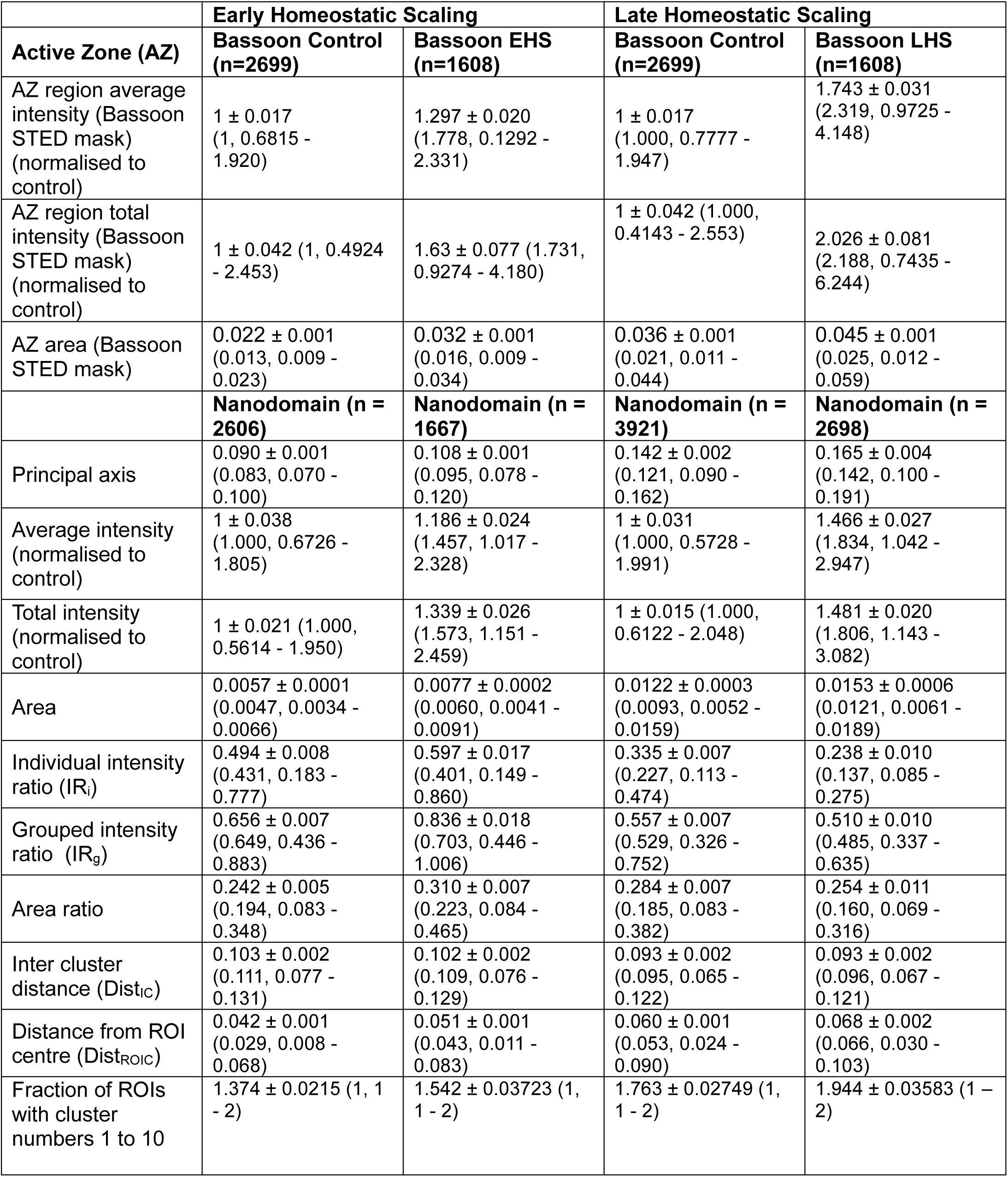
Data summarising the distribution of Bassoon in AZ during EHS and LHS.

**Table 3:** Data summarising the distribution of VGCC in synapses during EHS and LHS.

|  | **Early Homeostatic Scaling** |  | **Late Homeostatic Scaling** |  |
| --- | --- | --- | --- | --- |
| **Synaptic bouton** | **VGCC Control (n=267)** | **VGCC EHS (n=210)** | **VGCC Control (n=627)** | **VGCC LHS (n=372)** |
| Synaptic region average intensity (Bassoon confocal mask) (normalised to control) | $1 \pm 0.051$ (1, 0.5773 - 1.625) | $1.447 \pm 0.068$ (1.589, 0.9489 - 2.374) | $1 \pm 0.042$ (1.000, 0.5700 - 1.570) | $1.9 \pm 0.069$ (1.931, 1.195 - 2.969) |
| Synaptic region total intensity (Bassoon confocal mask) (normalised to control) | $1 \pm 0.093$ (1, 0.2507 - 2.937) | $1.49 \pm 0.151$ (1.424, 0.4619 - 4.1) | $1 \pm 0.058$ (1.000, 0.2825 - 2.825) | $2.52 \pm 0.174$ (2.092, 0.5403 - 8.753) |
| Synaptic area (Bassoon confocal mask) | $0.0677 \pm 0.0021$ (0.0428, 0.0164 - 0.0891) | $0.0784 \pm 0.0027$ (0.0502, 0.0203 - 0.1118) | $0.0857 \pm 0.0016$ (0.0576, 0.0239 - 0.1211) | $0.0992 \pm 0.0019$ (0.0719, 0.0286 - 0.1456) |
|  | **Nanodomain (n = 197)** | **Nanodomain (n = 170)** | **Nanodomain (n = 449)** | **Nanodomain (n = 227)** |
| Principal axis | $0.082 \pm 0.002$ (0.079, 0.069 - 0.093) | $0.080 \pm 0.002$ (0.077, 0.065 - 0.090) | $0.109 \pm 0.003$ (0.100, 0.078 - 0.121) | $0.138 \pm 0.007$ (0.116, 0.087 - 0.160) |
| Average intensity (normalised to control) | $1 \pm 0.076$ (1, 0.7667 - 1.408) | $0.983 \pm 0.074$ (1.071, 0.7870 - 1.435) | $1 \pm 0.059$ (1, 0.6531 - 1.575) | $1.601 \pm 0.141$ (1.448, 0.9217 - 2.997) |
| Total intensity (normalised to control) | $1 \pm 0.027$ (1, 0.7658 - 1.314) | $1.217 \pm 0.031$ (1.254, 1.002 - 1.637) | $1 \pm 0.044$ (1, 0.7631 - 1.337) | $1.629 \pm 0.053$ (1.784, 1.229 - 2.338) |
| Area | $0.0047 \pm 0.0002$ (0.0043, 0.0032 - 0.0060) | $0.0044 \pm 0.0002$ (0.0040, 0.0030 - 0.0054) | $0.0072 \pm 0.0003$ (0.0061, 0.0037 - 0.0091) | $0.0109 \pm 0.0008$ (0.0077, 0.0046 - 0.0141) |
| Individual intensity ratio ($IR_i$) | $0.342 \pm 0.024$ (0.281, 0.136 - 0.421) | $0.304 \pm 0.027$ (0.224, 0.137 - 0.353) | $0.212 \pm 0.009$ (0.155, 0.094 - 0.287) | $0.155 \pm 0.012$ (0.105, 0.068 - 0.185) |
| Grouped intensity ratio ($IR_g$) | $0.571 \pm 0.031$ (0.507, 0.277 - 0.890) | $0.554 \pm 0.033$ (0.532, 0.279 - 0.815) | $0.460 \pm 0.013$ (0.430, 0.289 - 0.616) | $0.284 \pm 0.014$ (0.234, 0.154 - 0.360) |
| Area ratio | $0.063 \pm 0.010$ (0.031, 0.017 - 0.063) | $0.056 \pm 0.013$ (0.026, 0.016 - 0.052) | $0.077 \pm 0.005$ (0.044, 0.022 - 0.093) | $0.089 \pm 0.013$ (0.050, 0.021 - 0.091) |
| Inter cluster distance ($Dist_{IC}$) | $0.296 \pm 0.013$ (0.295, 0.187 - 0.414) | $0.286 \pm 0.015$ (0.275, 0.178 - 0.399) | $0.219 \pm 0.006$ (0.201, 0.127 - 0.303) | $0.217 \pm 0.011$ (0.191, 0.110 - 0.324) |
| Distance from ROI centre ($Dist_{ROIc}$) | $0.154 \pm 0.022$ (0.144, 0.107 - 0.213) | $0.185 \pm 0.042$ (0.228, 0.086 - 0.263) | $0.141 \pm 0.005$ (0.126, 0.072 - 0.192) | $0.145 \pm 0.007$ (0.132, 0.078 - 0.202) |
| Fraction of ROIs with cluster numbers 1 to 10 | $2.079 \pm 0.1467$ (2, 1 - 2) | $2.152 \pm 0.1719$ (2, 1 - 3) | $2.038 \pm 0.1342$ (1, 1 - 2) | $2.124 \pm 0.2178$ (1, 1 - 2) |

**Table 4:** Data summarising the distribution of VGCC in AZ during EHS and LHS.

| Active Zone (AZ) | Early Homeostatic Scaling |  | Late Homeostatic Scaling |  |
| --- | --- | --- | --- | --- |
| AZ region average intensity (Bassoon STED mask) (normalised to control) | $1 \pm 0.037$<br>(1.000, 0.5042 - 1.695) | $1.325 \pm 0.036$<br>(1.473, 0.8668 - 2.310) | $1 \pm 0.030$<br>(1, 5023 - 1.785) | $1.755 \pm 0.045$<br>(1.919, 1.077 - 3.071) |
| AZ region total intensity (Bassoon STED mask) (normalised to control) | $1 \pm 0.066$ (1.000, 0.3840 - 3.051) | $1.657 \pm 0.129$<br>(1.641, 0.7131 - 4.338) | $1 \pm 0.051$ (1, 0.3258 - 2.802) | $2.502 \pm 0.151$<br>(1.869, 0.6661 - 7.629) |
| AZ area (Bassoon STED mask) | $0.0220 \pm 0.0006$<br>(0.0131, 0.0086 - 0.0225) | $0.0317 \pm 0.0013$<br>(0.0155, 0.0095 - 0.0338) | $0.0361 \pm 0.0007$<br>(0.0207, 0.0115 - 0.0439) | $0.0454 \pm 0.0011$<br>(0.0250, 0.0124 - 0.0590) |
|  | **Nanodomain (n = 226)** | **Nanodomain (n = 309)** | **Nanodomain (n = 727)** | **Nanodomain (n = 376)** |
| Principal axis | $0.081 \pm 0.003$ (0.077, 0.063 - 0.092) | $0.087 \pm 0.002$ (0.085, 0.069 - 0.099) | $0.110 \pm 0.003$ (0.098, 0.077 - 0.122) | $0.140 \pm 0.005$ (0.120, 0.085 - 0.158) |
| Average intensity (normalised to control) | $1 \pm 0.128$ (1, 0.7679 - 1.621) | $0.962 \pm 0.062$<br>(1.159, 0.7997 - 1.933) | $1 \pm 0.048$<br>(1, 0.6976 - 1.522) | $1.997 \pm 0.377$<br>(1.478, 0.9440 - 2.392) |
| Total intensity (normalised to control) | $1 \pm 0.026$ (1, 0.7805 - 1.279) | $1.365 \pm 0.036$<br>(1.353, 1.023 - 1.762) | $1 \pm 0.022$ (1, 0.7544 - 1.344) | $1.870 \pm 0.049$<br>(2.014, 1.291 - 2.578) |
| Area | $0.0045 \pm 0.0002$<br>(0.0041, 0.0027 - 0.0057) | $0.0052 \pm 0.0002$<br>(0.0049, 0.0031 - 0.0061) | $0.0075 \pm 0.0002$<br>(0.0062, 0.0038 - 0.0091) | $0.0115 \pm 0.0006$<br>(0.0087, 0.0046 - 0.0144) |
| Individual intensity ratio ($IR_i$) | $0.358 \pm 0.021$<br>(0.288, 0.138 - 0.552) | $0.346 \pm 0.020$<br>(0.225, 0.122 - 0.478) | $0.284 \pm 0.011$<br>(0.183, 0.091 - 0.377) | $0.200 \pm 0.013$<br>(0.119, 0.072 - 0.227) |
| Grouped intensity ratio ($IR_g$) | $0.501 \pm 0.018$<br>(0.434, 0.339 - 0.618) | $0.575 \pm 0.018$<br>(0.536, 0.333 - 0.772) | $0.515 \pm 0.011$<br>(0.464, 0.308 - 0.687) | $0.412 \pm 0.015$<br>(0.348, 0.249 - 0.501) |
| Area ratio | $0.114 \pm 0.012$<br>(0.057, 0.027 - 0.117) | $0.106 \pm 0.014$<br>(0.045, 0.023 - 0.103) | $0.146 \pm 0.008$<br>(0.083, 0.038 - 0.178) | $0.127 \pm 0.010$<br>(0.079, 0.034 - 0.149) |
| Inter cluster distance ($Dist_{IC}$) | $0.095 \pm 0.005$<br>(0.102, 0.071 - 0.129) | $0.100 \pm 0.003$<br>(0.107, 0.077 - 0.130) | $0.094 \pm 0.002$<br>(0.091, 0.068 - 0.122) | $0.073 \pm 0.005$<br>(0.075, 0.033 - 0.109) |
| Distance from ROI centre ($Dist_{ROIC}$) | $0.074 \pm 0.003$<br>(0.068, 0.044 - 0.100) | $0.079 \pm 0.003$<br>(0.080, 0.047 - 0.108) | $0.078 \pm 0.002$<br>(0.076, 0.049 - 0.107) | $0.084 \pm 0.003$<br>(0.080, 0.053 - 0.117) |
| Fraction of ROIs with cluster numbers 1 to 10 | $1.656 \pm 0.1166$ (1, 1 - 2) | $2.011 \pm 0.1351$ (1, 1 - 3) | $1.891 \pm 0.08741$ (1, 1 - 2) | $1.964 \pm 0.1219$ (1, 1 - 2) |

Structural parameters were direct entities extracted from a region of interest (such as a synapse or active zone) using morphometry analysis. The direct parameters provide information about the morphology and molecular content of the molecule of interest within a nanodomain, such as length, average intensity, and total intensity. In contrast, indirect or functional parameters provide information about the heterogeneity within the region of interest and require post hoc analysis, such as the fraction of molecules in nanodomains, total molecules, free molecules in a region, number of nanodomains within a region, and the distance between two nanodomains. The functional parameters we calculated by performing additional quantitative steps, like the Individual Intensity Ratio (IRi) (IR_i_ = total intensity of one nanodomain/total intensity of presynapse or active zone), Grouped Intensity Ratio (IRg) (IR_g_ = total intensity of all nanodomains/total intensity of presynapse or active zone), Area Ratio area of one nanodomain/area of presynapse or active zone), Inter-Cluster Distance (Dist_IC_) (distance between nanodomains), Distance from ROI Center (Dist_ROIC_) (distance of a nanodomain from the center of the region of interest (synapse or AZ)), Distance between Bassoon and VGCC Nanodomains (DistBV), and the number of nanodomains in each ROI. Though en extensive characterization of these organizational signature was lacking previously, our results are consistent with a few of the previous attempts ^8,19,30,34^. However, a holistic chareterization of the synaptic determinants that underlies lattice organization of molecules within and outside condensates reported exhaustively in Table 1-4 for basal conditions, EHS and LHS.

Since synaptic scaling affects several structural and functional parameters, we focused on those parameters that are altered differentially or antagonistically between EHS and LHS for both Bassoon and VGCC. When examining Bassoon during LHS, a reduction in IRi and IRg in AZ was observed (Fig. 4 A, B), implying a decrease in Bassoon within nanodomains in the AZ during LHS. During EHS, there was no effect on IRi in AZ, but IRg increased for Bassoon (Fig. 4 A, B). This data indicated that there was an increase in the clustering of Bassoon into nanodomains in the AZ during EHS, whereas an opposite effect was observed during LHS. For VGCC, a reduction in both IRi and IRg was observed in the AZ during LHS (Fig. 4 C, D). However, no deviation of IRi for VGCC was observed in EHS. Nevertheless, a small yet significant increase in IRg was observed (Fig. 4 C, D), implying that more VGCC molecules got clustered into nanodomains in a given AZ during EHS. This indicated an increase in free VGCC molecules at AZ, consistent with the phase transition data. During LHS, there was an increase in free energy for VGCC (Fig. 1 I), consistent with the occurrence of free VGCC molecules. Overall, dSTORM and STED data pointed towards increased free molecules for VGCC during LHS in the AZ. These observations were consistent with the phase transition data, which showed that EHS reduces the nucleation barrier, making it easier for VGCC to form clusters and not stay as free molecules (Fig. 1 F). However, when Bassoon was analyzed in the synapse, EHS had no effect (Fig. S12 A-B). In contrast, LHS showed an increase in free molecules with a reduction in both IRi and IRg (Fig. S12 A-B) consistent with previous reports ^34^. Similarly, in synapses, EHS did not have any effect on the number of free molecules of VGCC, while LHS showed an increase in free molecules with a reduction in both IRi and IRg (Fig S12 C-D). These observations indicate that the effects of homeostatic scaling are dependent on the functional zone, showing divergent patterns based on the specific protein and its localization and function within the synaptic and subsynaptic compartments.

**Figure 4:**
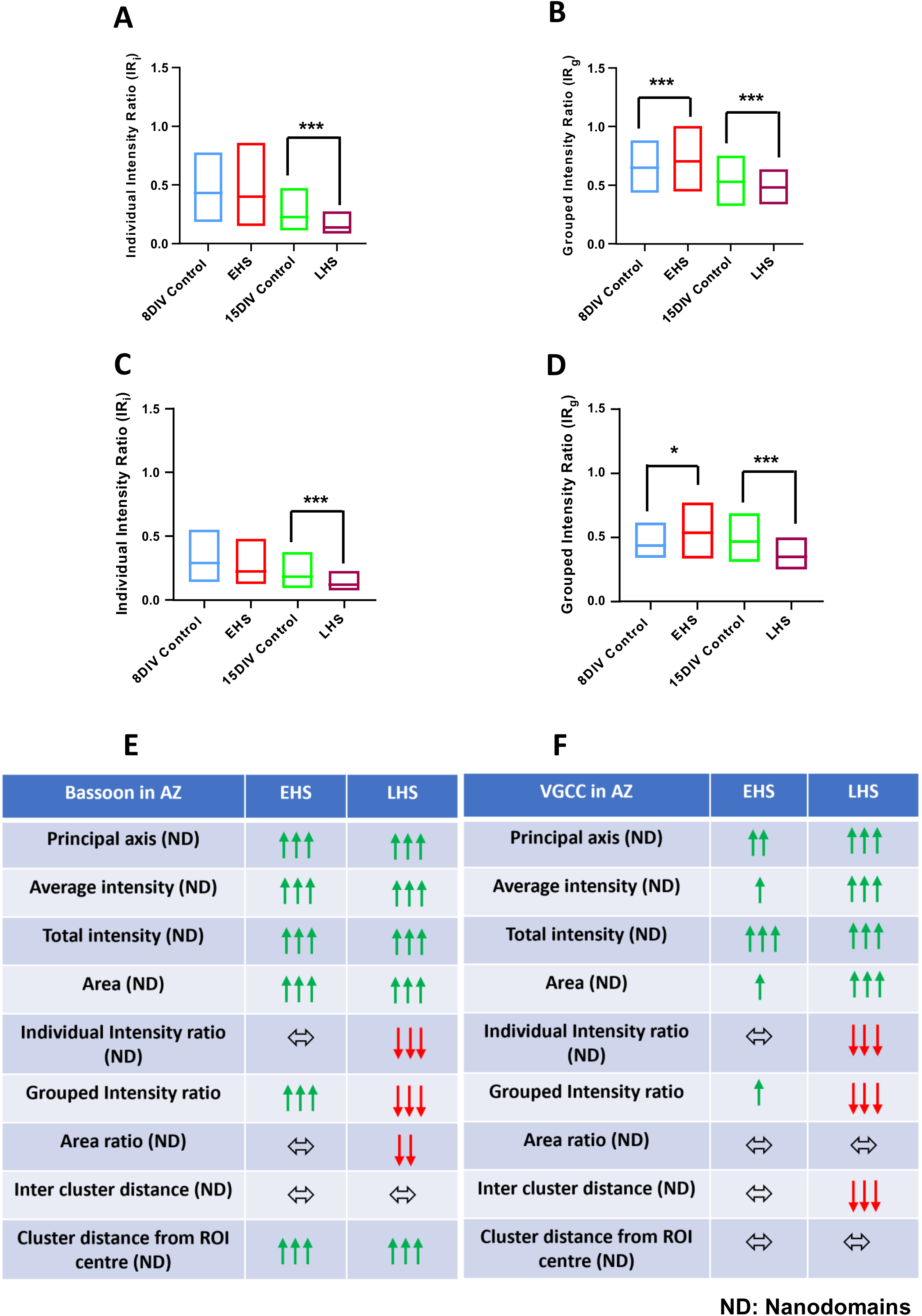
Effect of homeostatic scaling on Bassoon and VGCC in active zones (AZ): (A) Individual intensity ratio (IRi) of Bassoon nanodomains per AZ at DIV 8 between control and early homeostatic scaling (EHS) conditions (n=1686 and 1667 nanodomains for control and EHS datasets, respectively). Individual intensity ratio (IRi) of Bassoon nanodomains per AZ at DIV 15 between control and late homeostatic scaling (LHS) conditions (n=1640 and 688 nanodomains for control and LHS datasets, respectively). (B) Grouped intensity ratio (IRg) of Bassoon nanodomains per AZ at DIV 8 between control and EHS conditions (n=1686 and 1667 for control and EHS datasets, respectively). Grouped intensity ratio (IRg) of Bassoon nanodomains per synapse at DIV 15 between control and LHS conditions (n=1640 and 688 for control and LHS datasets, respectively). (C) Individual intensity ratio (IRi) of VGCC nanodomains per AZ at DIV 8 between control and EHS conditions (n=169 and 230 nanodomains for control and EHS datasets, respectively). Individual intensity ratio (IRi) of VGCC nanodomains per AZ at DIV 15 between control and LHS conditions (n=597 and 276 nanodomains for control and LHS datasets, respectively). (D) Grouped intensity ratio (IRg) of VGCC nanodomains per AZ at DIV 8 between control and EHS conditions (n=169 and 230 for control and EHS datasets, respectively). Grouped intensity ratio (IRg) of VGCC nanodomains per AZ at DIV 15 between control and LHS conditions (n=597 and 276 for control and LHS datasets, respectively). (E) Table representing the effect of homeostatic scaling on various structural and functional parameters of Bassoon in AZ. (F) Table representing the effect of homeostatic scaling on various structural and functional parameters of VGCC in AZ.

Upon closer examination of the structural parameters, it was observed that the principal axis and area of VGCC nanodomains remained unaffected in the synapse during EHS (Fig. S13 A-B). Additionally, the spatial distribution of VGCC nanodomains remained unchanged in both the synapse and AZ during EHS, as indicated by the lack of changes in Dist_IC_ and Dist_ROIC_ (Fig. S13 C, D, E and H). However, EHS affected VGCC nanodomains in the AZ, as evidenced by an increase in the principal axis (Fig. S13 E, DIV 8) and area of nanodomains (Fig. S13 F). On the contrary, LHS increased the principal axis and area of VGCC nanodomains in both the synapse and AZ (Fig. S.13 A, B, E and F). Notably, during LHS, the VGCC cluster distance from each other remained unchanged in synapses (Fig. S13 C), while there was a reduction in Dist_IC_ in the AZ (Fig. S.13 G, DIV 15), suggesting that the VGCC nanodomains moved closer to each other in the AZ. However, the Dist_ROIC_ stayed unaffected in both synapses (Fig. S13 D) and AZ (Fig. S13 H). In summary, LHS had a more potent effect on VGCC nanodomains by increasing their sizes in both synapses and AZ and causing them to move closer to each other in the AZ. When examining the structural parameters of Bassoon nanodomains, it was observed that the principal axis and area of nanodomains increased in both synapses and AZ during both EHS and LHS (Fig. S14A) However, during EHS, the Dist_IC_ of Bassoon reduced in synapses (Fig. S14 C), whereas the Dist_ROIC_ remained unchanged (Fig. S14 D). In contrast, the Bassoon Dist_IC_ remained unaffected in the AZ (Fig. S14G), but there was an increase in Dist_ROIC_ (Fig. S14 H). Essentially, homeostatic synaptic scaling caused the Bassoon nanodomains to move closer in synapses and spread out in the AZ during EHS. During LHS, the principal axis and area of Bassoon nanodomains increased in both synapses and AZ (Fig. S14 A,B). However, LHS reduced the Dist_IC_ of Bassoon nanodomains in synapses (Fig. S14 C) but not in the AZ (Fig. S14 G). Furthermore, there was an increase in Dist_ROIC_ of Bassoon nanodomains in the AZ during LHS (Fig. S14 H), indicating that the Bassoon nanodomains tended to move closer in synapses but spread out from the center of the AZ during both EHS and LHS.

### Subtle alterations in the nanoscale lattice arrangement of VGCC nanodomains and its proximity to AZ is sufficient to increase efficacy of neurotransmitter release

Two key observations of all the analysis were the altered molecular content of VGCC in nanodomains resulting from EHS and LHS, and the discrete distribution of VGCC with respect to Bassoon, which was used to mark AZ. We investigated using a data driven model of synaptic transmission obtained from our experiments to evaluate if subtle changes in spatial basis functions can fine tune probability of vesicle release at the active zone. As the localization of VGCCs and their proximity to docked vesicles can influence the probability of synaptic vesicle release, we relied on a nanoscale biophysical, stochastic, and spatially explicit model of the CA3 presynapse to simulate the calcium dynamics leading to vesicular release during action potential arrival ^8,18,35^. A nanoscale functional model of vesicular release was constructed using the relevant molecules in their physiological arrangements and concentrations consistent with our observation in a simplified cuboidal geometry with dimensions representing a canonical presynaptic compartment. Vesicle releases were simulated under different localization and assemblies of VGCC molecules to observe heterogeneity across the release events. (Fig 5A, B). The model incorporated vesicular release in all known pathways: calcium-dependent synchronous and asynchronous release, as well as calcium-independent spontaneous release^5,35,36^. VGCCs were positioned within nanodomains with a radius of 50 nm for all simulations, and the number of channels and the distance of the nanodomain from the release site were varied (Fig 5A, C) We evaluated the kinetics of release probability by placing the center of VGCC condensates 10 to 400 nm away from the release site. We also varied the number of VGCCs in the nanodomains from 2 to 200 molecules of VGCC for all coupling distances (Fig. 5C, D). The release probability showed a sigmoidal growth with the VGCCs number, irrespective of the coupling distance, and exhibited a steeper curve for closer distances. Next, we evaluated the effect on release probability by altering the content of VGCC in nanodomains akin to multiplicative scaling in Homeostatic plasticity (Fig 5E, F). For this, we kept every parameter the same but varied the scaling factors for synaptic configurations with either up or downscaling of molecular content within nanodomains (Fig. 5B, E, F). We found that over different distance ranges, the upscaling increased the probability of release events compared to unscaled conditions. On the contrary, downscaling decreased the release probability (Fig. 5E, F). This shows that subtle alteration of VGCC content is sufficient to have an agonistic or antagonistic outcome toward the probability of release.

**Figure 5:**
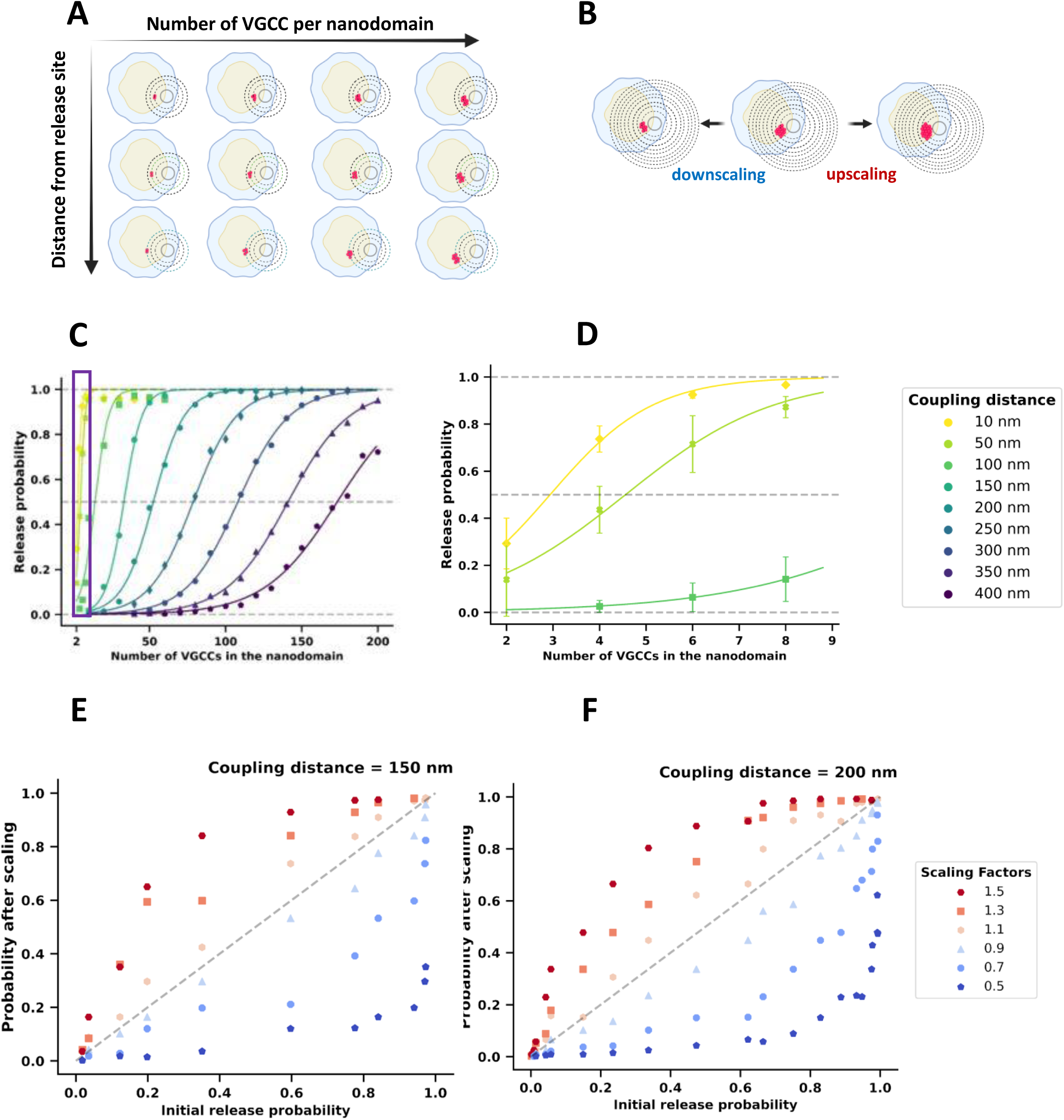
The molecular content of VGCC in nanocondensates and its proximity to active zones (AZ) modulates the efficacy of neurotransmitter release: (A) A schematic depicting the increase in the number of VGCC molecules per nanodomain from left to right. The top-down representation illustrates the increasing distance of the VGCC nanodomains from the neurotransmitter release site in the AZ. (B) A schematic representation of the upscaling and downscaling of VGCC in nanodomains. (C) The increase in the number of VGCC molecules (ranging from 10 to 200) is depicted on the x-axis, while the probability of release is shown on the y-axis. The different colored curves represent varying distances from the neurotransmitter release site (ranging from 10 nm to 400 nm). (D) This panel presents a zoomed-in view of the content of VGCC nanodomains, ranging from 2 to 8, at distances of 10 nm, 50 nm, and 100 nm from the neurotransmitter release site, as indicated by the red box in panel (C). The inset next to panel (D) shows the various distances considered for analysis, ranging from 10 nm up to 400 nm. (E-F) These panels illustrate the effect of different VGCC scaling factors on the probability of neurotransmitter release.(E) This panel depicts the effect of VGCC scaling factors ranging from 0.5 to 1.5 on the probability of release, operating at a distance of 150 nm from the neurotransmitter release site. (F) This panel illustrates the effect of VGCC scaling factors ranging from 0.5 to 1.5 on the probability of release, operating at a distance of 200 nm from the neurotransmitter release site. The inset next to panel (F) presents the exact scaling factors used for this analysis.

## Discussion

Miniature Excitatory Postsynaptic Currents/Potentials (mEPSCs) are a consequence of spontaneous neurotransmitter release events, which play a crucial role in basal neuronal activity and synaptic maintenance ^5,15,36,37^. While several synaptic molecules are implicated in spontaneous release events, the exact mechanisms that modulate these events remain unclear^36^. A significant proportion of such events are modulated by VGCCs present in and around the AZ of the presynaptic compartment of an excitatory synapse^15^. Homeostatic scaling induced on younger neurons resulted in a change in amplitude while the older neurons show a change in frequency of miniature EPSCs^38^. Our observations confirms the recent evidence indicating the reorganization and reversible incomplete dissociation of the active zone matrix due to prolonged incubation with TTX, inducing homeostatic scaling ^39^. Our observations on lattice arrangement of VGCCs and Bassoon align with electrophysiological and ultrastructural findings, reinforcing the idea of differential regulation within the AZ. This evidence shows that synaptic transmission, which directly influences the characteristics of mEPSCs, is also modulated through distinct mechanisms at the molecular level within the active zone^5,18,27,40-42^.

Our observations revisit the molecular organization behind synaptic plasticity, a fundamental process in neural communication, through the lens of nanoscale molecular organization and homeostatic scaling. Using advanced methodologies like liquid-liquid phase separation (LLPS), microscopy and molecular modeling, we show how synapses adapt their structure and function to optimize information transfer during different stages of neuronal development using “compositional degeneracy” in EHS and LHS. We show that lateral arrangement of synaptic molecules is cardinal for compositional degeneracy, where synapses use varying molecular configurations to achieve the same functional outcomes, thus enhancing resilience and efficiency in information processing. Central to our investigation is the role of VGCC and Bassoon, a key structural protein at the active zone of synapses^16,34^. Research demonstrates that synapses undergo different thermodynamic changes in EHS and LHS. In EHS, molecules like VGCC and Bassoon cluster together, reducing free energy and promoting stability. Conversely, during LHS,these molecules behave more freely, increasing entropy. This phase-based organization highlights how synaptic compartments use thermodynamic properties to regulate excitability and information flow. The concept of synaptic plasticity as an information processing machine challenges traditional ideas of a one-to-one correlation between structural and functional changes. Instead, experimental observations coupled with molecular modelling verifies that synaptic computation results from complex, localized adjustments in nanoscale molecular structures, especially through the formation of nanodomains—tiny machines within synapses that govern molecular interactions^5,7,43,44^. These nanodomains of VGCC and Bassoon were mapped using multi-parameter image analysis, providing insights on how synapses control the probability of neurotransmitter release and adapt to changes in activity. Three key insights emerge from this study. First, it is crucial to consider thermodynamic uncertainty and information processing when analyzing synaptic plasticity. By examining relative changes in entropy and free energy, synapses optimize and fine-tune their information transfer capabilities. Second, this study addresses how these thermodynamic changes relate to plasticity induction—whether synapses enhance or reduce their efficacy by altering local precision by modulating entropy. Third, our results generates a detailed nanoscale caretesian map of molecular distribution, identifying how VGCC and Bassoon rearrange during different plasticity states to regulate neurotransmission. Thus, compositional degeneracy could provide multiple structural routes to preserve the signatures of synaptic transmission through the stochastic organization of VGCCs^9,45^.

The molecular organization underlying synaptic transmission is governed by two primary mechanisms. The first involves modulating synaptic strength through the addition of unitary synaptic complexes, while the second relies on spontaneous synaptic organization driven by liquid-liquid phase transitions^7,21,43^. These processes work in synchrony, with phase transition facilitating rapid, short-term reorganization of synaptic components aided by rapid exchange through diffusive processes, and the regulated addition of complexes with defined packing densities support long-term stability, which may require protein synthesis^5,19,29,46,47^. Additionally, such scaling is well correlated with the frequency and amplitude of synaptic transmission^12,22^. This study highlights that not all synaptic elements adhere to multiplicative scaling typically seen during homeostatic scaling^22,23^. For example, different scaffolding proteins form nanodomains with altered exchange rates or limited molecular numbers^1,6,13^. Our analysis further emphasizes the role of multiplicative scaling in synaptic plasticity. Voltage-gated calcium channels exhibit distinct scaling in different synaptic compartments^23^. Interestingly, while VGCC displays multiplicative scaling, Bassoon does not, highlighting the molecular heterogeneity within synapses. It could be speculated that different molecules have agonistic or antagonistic changes in entropy, preventing a complete breakdown of synaptic organization. This underscores the importance of nanoscale degeneracy, which enables diverse molecular configurations to maintain synaptic function by regulating transmission rates. We thus propose that synaptic transmission changes should be viewed as an optimization process, maximizing information transfer efficiency and ensuring functional robustness through its structural flexibility. The framework of nanoscale degeneracy provides synapses with the flexibility to select the most suitable state for a given condition, drawing from a wide range of structural assemblies on synaptic lamina to achieve similar functional outcomes. These inferences connect synaptic information transfer efficiency with probabilistic inferences, multiplicative changes in synaptic strength, and the spatial organization of structural markers such as Bassoon and functional markers like VGCC. Together, these factors create a nanoscale biochemical map that explains heterogeneity in synaptic transmission at the single-synapse level.

We illucidate that structural changes during homeostatic scaling are not uniform across molecules and compartments, yet collectively fine-tune synaptic function. We show that nanoscale compositional degeneracy at the level of individual synapses—where multiple molecular pathways can achieve the same functional outcomes—is key to optimizing synaptic transmission. This challenges the conventional view that structural reorganization directly correlates with function, instead revealing that computation emerges from subtle nanoscale adjustments rather than large-scale architectural shifts.This framework of compositional degeneracy allows synapses to remain functionally robust, ensuring efficient information transfer even with changes in structural configurations. By integrating thermodynamic principles, molecular heterogeneity and spatial organization, this research work offers a perspective on how synapses balance plasticity and stability in neural circuits. It also raises an age-old debate on whether we should revisit how we evaluate the quantal nature of synaptic transmission^48^ based on emerging information from a decade of nanoconnectomics data^5,37,49^.

## Acknowledgments

We extend our gratitude to all members of the Nano-organization Lab, Centre for Neuroscience, IISc, for their valuable discussions and feedback. We also thank Jean-Baptiste Sibarita, IINS, Bordeaux, for sharing analysis paradigms for single-molecule localization and tracking. This work was largely supported by grants awarded to D.N. from the DBT-Wellcome Trust India Alliance Senior Fellowship, the Indian Institute of Science under the Institute of Eminence (IISc-IOE) program, the Core Research Grant through ANRF, the Ignite Life Science Foundation, and the FABRIC program via the CBR-IISc partnership platform. P.R.N. received support from ICMR and IISc. We are also grateful for the support provided by the Central Animal Facility and Core Imaging Facility of IISc, the Microscopy Research Core of IISER Pune, and NCBS Bangalore.We also thank Prof. Rishikesh Narayanan, IISc for coining the term “Compositional Degeneracy”

## Author Contributions Statement-Inclusion and Ethics

D.N. conceived the project, established the necessary infrastructure, and conceptualized the data analysis pipelines and foundational imaging protocols. Together, P.R.N. and D.N. designed the research experiments and contributed to conceptual advances in experimental and analytical approaches. P.R.N. conducted all experiments and performed the data analysis. Y.C. developed data interfaces for extracting mathematical parameters, enabling further extensions of mathematical analysis. SN and RA conceptualized and developed the computational model for the presynaptic terminal and wrote the relevant methods and results for the simulations. Based on the processed data provided by DN and PRN, RA performed computational model simulations. S.N. provided guidance and interpretation of the synaptic transmission model. P.R. assisted with the analysis of phase transition dynamics, while S.K. supported P.R.N. in optimizing the initial data analysis. P.R.N. prepared the primary neuronal cultures, with M.J. assisting in their optimization, troubleshooting experiments, and co-editing the manuscript alongside P.R.N. and D.N. S.N. provided guidance and interpretation of the synaptic transmission model. P.R.N. and D.N. jointly wrote the manuscript. All authors reviewed the manuscript, offered critical input, and approved the final version.

## Supplementary Figure Legends

**Supplementary Figure 1: Scheme 1: Distribution of Bassoon and VGCC in a synapse.** The diagram on the top represents a presynaptic bouton that releases neurotransmitters encapsulated in a synaptic vesicle at the active zone (AZ). The diagram below is a magnified representation of a synapse where Bassoon and VGCC’s spatial distribution is depicted in the synapse and AZ. Bassoon and VGCC are represented as free molecules and clusters in the synapse and AZ.

**Supplementary Figure 2: HILO and dSTORM image of a young neuron with Bassoon and VGCC distribution in synapse and AZ.** Pseudocolor image of Bassoon and VGCC obtained using HILO microscopy (A and E) and dSTORM microscopy (C and G). Bassoon regions were automatically detected by thresholding the image (A) and are marked by black contours, which are representative of synaptic regions. The red contour within the black Bassoon confocal contour in C and G represents automatically detected dSTORM regions of Bassoon, representative of AZ region. Insets 1-5 represent potential synaptic regions with Bassoon distribution (A) Pseudocolor HILO image of Bassoon in a DIV 8 neuron. (B) HILO image of the DIV 8 neuron depicting the distribution of Bassoon in synapses. (C) dSTORM image of the DIV 8 neuron exhibiting the distribution of Bassoon in AZ. (D) dSTORM image of the DIV 8 neuron depicting the automatically detected clusters of Bassoon represented as black squares. (E) pseudocolor HILO image of VGCC in the DIV 8 neuron. Insets 6-10 represent potential synaptic regions with VGCC distribution (F) HILO image of the DIV 8 neuron displaying the distribution of VGCC in synapses. (G) dSTORM image of the DIV 8 neuron depicting the distribution of VGCC in AZ. (H) dSTORM image of the DIV 8 neuron displaying the automatically detected clusters of Bassoon and VGCC represented as green and red squares, respectively. Scale bar in H is 10 microns.

**Supplementary Figure 3: HILO and dSTORM image of a DIV 15 neuron with Bassoon and VGCC distribution in synapse and AZ.** Pseudocolor image of Bassoon and VGCC obtained using HILO microscopy (A and E) and dSTORM microscopy (C and G). Bassoon regions were automatically detected by thresholding the image (A) and are marked by black contours, which are representative of synaptic regions. The red contour within the black Bassoon confocal contour in C and G represents automatically detected dSTORM regions of Bassoon, representative of AZ region. (A) Pseudocolor HILO image of Bassoon in a DIV 15 neuron. Insets 1-5 represent potential synaptic regions with Bassoon distribution (B) HILO image of the DIV 15 neuron depicting the distribution of Bassoon in synapses. (C) dSTORM image of the DIV 15 neuron depicting the distribution of Bassoon in AZ. (D) dSTORM image of the DIV 15 neuron depicting the automatically detected clusters of Bassoon represented as black squares. (E) Pseudocolor HILO image of VGCC in the DIV 15 neuron. Insets 6-10 represent potential synaptic regions with VGCC distribution (F) HILO image of the DIV 15 neuron depicting the distribution of VGCC in synapses. (G) dSTORM image of the DIV 15 neuron depicting the distribution of VGCC in AZ. (D) dSTORM image of the DIV 15 neuron depicting the automatically detected clusters of Bassoon and VGCC represented as green and red squares, respectively. Scale bar in H is 10 microns.

**Supplementary Figure 4: Scheme 2 - Phase transition of Bassoon and VGCC. (**A) The diagram above is a visual representation of aggregates of single molecules undergoing a first-order phase transition. (B) A plot of the probability distribution function of single molecules detected inside the nanoclusters. (C) The curve fit of the inverse of the probability distribution of molecules according to the function an^2/3^–bn+c to obtain the parameters a, b, and c, which defines the nucleation barrier (ΔGc) and critical cluster radius (Rc). (D) Linear regression plot of the resultant surface energy correction by subtracting the surface energy (an^2/3^) from the –LogP(n) data. The negative linear slope indicates the second term, b, which corresponds to an increase in entropy. (E) Schematic of the free energy function, which follows the first-order phase transition.

**Supplementary Figure 5: Effect of early and late homeostatic scaling on the slope of bulk energy parameter-bn**. (A-H) Linear regression plot of the resultant of surface energy correction by subtracting the n^2/3^ surface energy term from the Log P(n) data. The negative linear slope (λ) suggests the second term bn. The effect of homeostatic scaling on the slope of bn in (A) DIV 8 Bassoon control, (B) DIV 8 Bassoon EHS (C) DIV 8 VGCC control (D) DIV 8 VGCC EHS (E) DIV 15 Bassoon control (F) DIV 15 Bassoon LHS (G) DIV 15 VGCC control (H) DIV 15 VGCC LHS.

**Supplementary Figure 6: Scheme 3 - Multiplicative scaling analysis** (A) The diagram above represents the spatial distribution of Bassoon and VGCC inside the synapse and AZ. For the ease of explaining the analysis protocol, the distribution of Bassoon and VGCC have been split into red for VGCC and green for Bassoon. The protocol is described for the multiplicative scaling induced for VGCC in the synapse. Briefly, (B) the histogram represents the average intensity of VGCC inside the synapse. The intensity values were rank ordered separately for control and EHS/LHS conditions and the straight line (C) was fit to the resulting data. After, the EHS/LHS dataset was scaled by multiple arbitrary scaling factors (SF); the SF corresponding to the highest p-value was considered the most accurate. K-S test was used to compare between control and scaled distributions. This entire process was repeated 100 times from random sampling of control and EHS/LHS to get an average SF (D). Once the desired SF was obtained, the EHS/LHS dataset was divided by the SF and the resulting distribution was termed as scaled distribution. (E) Finally, the cumulative frequency plot of control, EHS/LHS (TTX) and scaled TTX distributions was plotted, showing successful multiplicative scaling.

**Supplementary Figure 7: Linear fit of control and TTX datasets for average intensity of VGCC in synapses and AZ during EHS and LHS.** From both control and TTX data, the same number of puncta were chosen randomly. EHS/LHS (TTX) data were plotted against the control data, and a linear fit was calculated after rank ordering from lowest to highest fluorescence intensity. An estimate for the multiplicative scaling factor was given by the slope of the linear fit. (A-D) Linear fit for VGCC average intensity in synapses and AZ during EHS (A, B respectively) (n=267 and 210 for control and EHS dataset in synapses; n=703 and 601 for control and EHS dataset in AZ) and LHS (C, D respectively) (n=627 and 372 for control and LHS dataset in synapses; n=1386 and 825 for control and LHS dataset in AZ).

**Supplementary Figure 8: Effect of early homeostatic scaling on the average intensity of Bassoon in synapses and active zones (AZ).** (A) Representation of the distribution of Bassoon average intensity per synapse in a DIV 8 neuron in control and EHS conditions (n=1175 and 873 synapses for control and EHS dataset). (B) Multiplicative scaling factor for Bassoon in the synapse at DIV 8 (1.24, p= 3.85e-12, K-S test). (C) Normalized cumulative frequency distribution of control, TTX and scaled TTX for Bassoon average intensity in synapses at DIV 8. (D) Representation of the distribution of Bassoon average intensity per AZ in a young neuron in control and the TTX-treated condition (n=2699 and 1608 AZ for control and TTX dataset). (E) Multiplicative scaling factor for Bassoon in the AZ at DIV 8 (1.1, p= 3.71e-10, K-S test) F) Normalized cumulative frequency distribution of control, TTX and scaled TTX for Bassoon average intensity in AZ at DIV 8. (G) Representation of the distribution of average intensity of Bassoon nanodomains per synapse in a DIV 8 neuron in control and EHS conditions (n=1347 and 898 synapses for control and EHS dataset). (H) Multiplicative scaling factor for Bassoon in the synapse at DIV 8 (1.1, p= 0.00458, K-S test). I) Normalized cumulative frequency distribution of control, TTX and scaled TTX for Bassoon nanodomain average intensity in synapses at DIV 8. (J) Representation of distribution of Bassoon nanodomain average intensity per AZ in a young neuron in control and the TTX treated conditions (n=2606 and 1667 AZ for control and TTX dataset). (K) Multiplicative scaling factor for Bassoon nanodomains in the AZ at DIV 8 (1.12, p= 0.00796, K-S test). L) Normalized cumulative frequency distribution of control, TTX and scaled TTX for Bassoon nanodomain average intensity in AZ at DIV 8.

**Supplementary Figure 9: Effect of late homeostatic scaling on Bassoon average intensity in synapses and active zones (AZ)** (A) Representation of the distribution of Bassoon average intensity per synapse in a DIV 15 neuron in control and LHS conditions (n=2567 and 2224 synapses for control and LHS dataset). (B) Multiplicative scaling factor for Bassoon in the synapse at DIV 8 (1.73, p= 1.47e-5, K-S test). C) Normalized cumulative frequency distribution of control, TTX and scaled TTX for Bassoon average intensity in synapses at DIV 15. (D) Representation of distribution of Bassoon average intensity per AZ in a young neuron in control and the TTX-treated conditions (n=3481 and 2523 AZ for control and TTX dataset). (E) Multiplicative scaling factor for Bassoon in the AZ at DIV 15 (2.64, p= 4.68e-132, K-S test). F) Normalized cumulative frequency distribution of control, TTX and scaled TTX for Bassoon average intensity in AZ at DIV 15. (G) Representation of the distribution of the average intensity of Bassoon nanodomains per synapse in a DIV 15 neuron in control and LHS conditions (n=2830 and 2532 synapses for control and EHS dataset). (H) Multiplicative scaling factor for Bassoon in the synapse at DIV 15 (1.42, p= 1.7e-5, K-S test). I) Normalized cumulative frequency distribution of control, TTX and scaled TTX for Bassoon nanodomain average intensity in synapses at DIV 15. (J) Representation of the distribution of Bassoon nanodomain average intensity per AZ in a young neuron in control and the TTX-treated conditions (n=3921 and 2698 AZ for control and TTX dataset). (K) Multiplicative scaling factor for Bassoon nanodomains in the AZ at DIV 15 (1.96, p= 0.0011, K-S test). L) Normalized cumulative frequency distribution of control, TTX and scaled TTX for Bassoon nanodomain average intensity in AZ at DIV 15.

**Supplementary Figure 10: Parity between dSTORM and STED data for Bassoon and VGCC.** The data from dSTORM and STED were compared to understand if the different microscopy techniques changed the observations. (A) The number of molecules of Bassoon as obtained from dSTORM in synapses at DIV 15 in control (N_ND_Syn_Con_) and after LHS (N_ND_Syn_LHS_). (B) The number of molecules of VGCC as obtained from dSTORM in synapses at DIV 15 in control (N_ND_Syn_Con_) and after LHS (N_ND_Syn_LHS_). (C) The number of molecules of Bassoon as obtained from dSTORM in AZ at DIV 15 in control (N_ND_AZ_Con_) and after LHS (N_ND_AZ_LHS_). (D) The number of molecules of VGCC as obtained from dSTORM in AZ at DIV 15 in control (N_ND_AZ_Con_) and after LHS (N_ND_AZ_LHS_). (E) The total intensity of Bassoon nanodomains in synapses at DIV 15 as obtained from STED in control and after LHS. (F) The total intensity of VGCC nanodomains in synapses at DIV 15 as obtained from STED in control and after LHS. (G) The total intensity of Bassoon nanodomains in AZ at DIV 15 as obtained from STED in control and after LHS. (H) The total intensity of VGCC nanodomains in AZ at DIV 15 as obtained from STED in control and after LHS. The n values for the number of molecules of Bassoon in nanodomains in synapses and AZ were control: 2301, LHS: 2364 and for VGCC control: 1270, LHS: 1892. The n values for the total intensity of Bassoon in synaptic nanodomains at DIV 15 control: 1106, LHS: 620 and for VGCC, control: 382, LHS:151. The n values for the total intensity of Bassoon in nanodomains in AZ DIV 15 control: 1640, LHS: 688 and for VGCC, control: 597, LHS: 276.

**Supplementary Figure 11: Scheme 4 - Parameters extracted from nanodomains obtained via STED microscopy for analyzing the distribution of Bassoon and VGCC in different compartments**. Different regions of interest (ROIs) were analyzed differentially. For analysing synapse as an ROI, a confocal mask of Bassoon was utilized. For analysing the AZ as an ROI, STED mask of Bassoon was used, and finally, automatically detected clusters were used as the mask for nanodomains. Different structural and functional parameters were extracted from the data obtained from STED microscopy. Direct parameters include Intensity of Bassoon/VGCC within a synapse, AZ and nanodomain: Total intensity (sum of all grayscale values of all pixels within a given ROI) and Average intensity (Total intensity/number of pixels within that ROI), principal axis of a nanodomain and area of a nanodomain. Specific functional parameters were extracted, such as Individual intensity ratio (IR_i_) = total intensity of one nanodomain/total intensity of presynapse or active zone, Grouped intensity ratio (IR_g_) = total intensity of all nanodomains/total intensity of presynapse or active zone, Area ratio = Area of one nanodomain/Area of presynapse. Spatial parameters were also extracted such as Inter cluster distance (Dist_IC_) which measures the distance between two nanodomains; distance from ROI (region of interest) center (Dist_ROIC_) was calculated as the distance of a nanodomain from the center of the ROI, Intercluster distance between Bassoon and VGCC (Dist_IC_BV) where distances between Bassoon and VGCC within a single synapse were calculated. Information was extracted from ROIs containing 1-10 cluster numbers.

**Supplementary Figure 12: Effect of homeostatic scaling on the amount of free Bassoon and VGCC molecules in synapse.** (A) Individual intensity ratio (IRi) of Bassoon nanodomain per synapse at DIV 8 between the control and the EHS conditions (n= 809 and 776 nanodomains for control and EHS dataset). Individual intensity ratio (IRi) of Bassoon nanodomain per synapse at DIV 15 between the control and the LHS conditions (n=1106 and 620 nanodomains for control and LHS dataset). (B) Grouped intensity ratio (IRg) of Bassoon nanodomain per synapse at DIV 8 between the control and the EHS conditions (n=809 and 776 for control and TTX dataset). Grouped intensity ratio (IRg) of Bassoon nanodomain per synapse at DIV 15 between the control and the LHS conditions (n=1106 and 620 for control and LHS dataset). (C) Individual intensity ratio (IRi) of VGCC nanodomain per synapse at DIV 8 between the control and the EHS conditions (n=128 and 97 nanodomains for control and EHS dataset). Individual intensity ratio (IRi) of VGCC nanodomain per synapse at DIV 15 between the control and the LHS conditions (n= 382 and 151 nanodomains for control and LHS dataset). (D) Grouped intensity ratio (IRg) of VGCC nanodomain per synapse at 8 DIV between the control and the EHS conditions (n=128 and 97 for control and EHS dataset). Grouped intensity ratio (IRg) of VGCC nanodomain per synapse at DIV 15 between the control and the LHS conditions (n= 382 and 151 for control and LHS dataset).

**Supplementary Figure 13: Effect of homeostatic scaling on the dimensional and spatial parameters of VGCC in the synapse and AZ of a neuron**. (A) Principal axis of VGCC nanodomain per synapse at DIV 8 between the control and the EHS conditions (n=128 and 97 nanodomains for control and EHS dataset). Principal axis of VGCC nanodomain per synapse at DIV 15 between the control and the LHS conditions (n=382 and 151 nanodomains for control and LHS dataset). (B) Area of VGCC nanodomain per synapse at DIV 8 between the control and the EHS conditions (n= 128 and 97 nanodomains for control and EHS dataset). Area of VGCC nanodomain per synapse at DIV 15 between the control and the LHS conditions (n= 382 and 151 nanodomains for control and LHS dataset). (C) Inter cluster distance (Dist_IC_) between VGCC nanodomains per synapse at DIV 8 between the control and the EHS conditions (n= 87 and 71 values for control and EHS dataset). Inter cluster distance (Dist_IC_) between VGCC nanodomains per synapse at DIV 15 between the control and the LHS conditions (n= 388 and 107 values for control and LHS dataset). (D) Distance from ROI centre (Dist_ROIC_) of VGCC nanodomains per synapse at DIV 8 between the control and the EHS conditions (n= 160 and 181 values for control and EHS dataset). Distance from ROI centre (Dist_ROIC_) of VGCC nanodomains per synapse at DIV 15 between the control and the LHS conditions (n= 382 and 151 values for control and LHS dataset). (E) Principal axis of VGCC nanodomain per AZ at DIV 8 between the control and the EHS conditions (n= 169 and 230 nanodomains for control and EHS dataset). Principal axis of VGCC nanodomain per AZ at DIV 15 between the control and the LHS conditions (n=597 and 276 nanodomains for control and LHS dataset). (F) Area of VGCC nanodomain per AZ at DIV 8 between the control and the EHS conditions (n= 169 and 230 nanodomains for control and TTX dataset). Area of VGCC nanodomain per AZ at DIV 15 between the control and the LHS conditions (n= 597 and 276 nanodomains for control and LHS dataset). (G) Inter cluster distance (Dist_IC_) between VGCC nanodomains per AZ at DIV 8 between the control and the EHS conditions (n=59 and 107 values for control and EHS dataset). Inter cluster distance (Dist_IC_) between VGCC nanodomains per AZ at DIV 15 between the control and the LHS conditions (n=193 and 113 values for control and LHS dataset). (H) Distance from ROI centre (Dist_ROIC_) of VGCC nanodomains per AZ at DIV 8 between the control and the EHS conditions (n=160 and 181 values for control and TTX dataset). Distance from ROI centre (Dist_ROIC_) of VGCC nanodomains per AZ at DIV 15 between the control and the LHS conditions (n= 420 and 169 values for control and TTX dataset).

**Supplementary Figure 14: Effect of homeostatic scaling on the dimensional and spatial parameters of Bassoon nanodomains in synapse and AZ of a neuron.** (A) Principal axis of Bassoon nanodomain per synapse at DIV 8 between the control and the EHS conditions (n= 809 and 776 nanodomains for control and EHS dataset). Principal axis of Bassoon nanodomain per synapse at DIV 15 between the control and the LHS conditions (n=1106 and 620 nanodomains for control and LHS dataset). (B) Area of Bassoon nanodomain per synapse at DIV 8 between the control and the EHS conditions (n= 809 and 776 nanodomains for control and EHS dataset). Area of Bassoon nanodomain per synapse at DIV 15 between the control and the LHS conditions (n=1106 and 620 nanodomains for control and LHS dataset). (C) Inter cluster distance (Dist_IC_) between Bassoon nanodomains per synapse at DIV 8 between the control and the EHS conditions (n=682 and 631 values for control and EHS dataset). Inter cluster distance (Dist_IC_) between Bassoon nanodomains per synapse at DIV 15 between the control and the LHS conditions (n=944 and 652 values for control and LHS dataset). (D) Distance from ROI centre (Dist_ROIC_) of Bassoon nanodomains per synapse at DIV 8 between the control and the EHS conditions (n=672 and 686 values for control and EHS dataset). Distance from ROI centre (Dist_ROIC_) of Bassoon nanodomains per synapse at DIV 15 between the control and the LHS conditions (n=1106 and 620 values for control and LHS dataset). (E) Principal axis of Bassoon nanodomain per AZ at DIV 8 between the control and the EHS conditions (n=1686 and 1667 nanodomains for control and EHS dataset). Principal axis of Bassoon nanodomain per AZ at DIV 15 between the control and the LHS conditions (n= 1640 and 688 nanodomains for control and LHS dataset). (F) Area of Bassoon nanodomain per AZ at DIV 8 between the control and the EHS conditions (n= 1686 and 1667 nanodomains for control and EHS dataset). Area of Bassoon nanodomain per AZ at DIV 15 between the control and the LHS conditions (n= 1640 and 688 nanodomains for control and TTX dataset). (G) Inter cluster distance (Dist_IC_) between Bassoon nanodomains per AZ at DIV 8 between the control and the EHS conditions (n=329 and 347 values for control and EHS dataset). Inter cluster distance (Dist_IC_) between Bassoon nanodomains per AZ at DIV 15 between the control and the LHS conditions (n=489 and 316 values for control and LHS dataset). (H) Distance from ROI centre (Dist_ROIC_) of Bassoon nanodomains per AZ at DIV 8 between the control and the EHS conditions (n=1470 and 1188 values for control and EHS dataset). Distance from ROI centre (Dist_ROIC_) of Bassoon nanodomains per AZ at DIV 15 between the control and the LHS conditions (n=1285 and 501 values for control and LHS dataset).

## Data Availability Statement

Example dataset are available for download on Github - https://github.com/Yuktichopra/vgcc-bassoon in the raw_files of data_and_code section.

## Code availability statement

The presented work utilized already available codes.

The version of the code used for extraction of data for multiplicative scaling and multi parameter analysis use are available for download on Github https://github.com/Yuktichopra/vgcc-bassoon along with example extracted parameters and scripts. The set of instructions to use the analysis modules are explained in the supplementary information uploaded in the folder.

PALMTracer is an all-in-one software package for the analysis of Single Molecule Localization Microscopy (SMLM) data which can be downloaded from https://neuro-intramuros.u-bordeaux.fr/displayresearchprojects/70/11

## Methods

### Materials and Methods

#### Experimental animals

Wild-type (WT) Sprague-Dawley rat pups were acquired from the Central Animal Facility, IISc. These animals were bred and housed in a pathogen-free environment which was temperature controlled in a 12 h light/12 h dark cycle. The animals had ad libitum access to food and water. All animal experiments were performed under strict guidelines and approval from the Institutional Animal Ethics Committee (IAEC), Central Animal Facility (CAF), Indian Institute of Science, Bangalore, India ^1-3^

#### Primary hippocampal cultures

Hippocampal neurons were extracted from postnatal day 0–1 Sprague Dawley rat pups. The dissected hippocampi were trypsinized (Thermo Fisher Scientific) and dissociated in Hibernate-A (Thermo Fisher Scientific) media supplemented with B27 (Thermo Fisher Scientific), GlutaMAX (Thermo Fisher Scientific), and 100 mg/ml Normocin (Invivogen). Once dissociated, neurons were resuspended in Neurobasal A (Thermo Fisher Scientific) medium supplemented with B27, GlutaMAX and 100 mg/ml Normocin (InvivoGen). The cells were seeded in 12 well plates at a density of 0.1×10^6^ cells/mL in 18 mm #1.5 (corrected for 0.17+/- 0.01) glass coverslips coated with poly-L-lysine at a concentration of 100 µg/mL. Fresh Neurobasal A medium was supplemented every week to maintain the viability of the culture^2,4^

#### Homeostatic plasticity induction protocol

Induction of homeostatic scaling was conducted by incubation of rat hippocampal neuronal cultures with 2 µM TTX (Alamone labs) for 24 h either at 7 days in vitro (DIV 7) or at DIV 14. The cultures treated with TTX were referred to as TTX dataset, whereas untreated cultures of corresponding age were used as control ^2,5^

#### Immunocytochemistry

The hippocampal neuronal cells were fixed by incubating with 4% paraformaldehyde supplmented with 4% sucrose in PBS at 4°C for 10 min. The cells were then quenched with 0.1 M glycine in PBS at room temperature, permeabilized with 0.25% Triton X-100 for 5 minutes and then blocked with 10% BSA in PBS for 30 minutes at room temperature. The primary antibody was incubated on the cells for 1–2 h and then washed four times (5 minutes each) with 3% BSA. After,the cells were incubated with a suitable secondary antibody for 45 minutes and then washed four times with PBS (5 minutes each). The cells were mounted with Prolong (Molecular Probes, cat. no. MAN0010261) for visualization under confocal/STED imaging^4,6^.

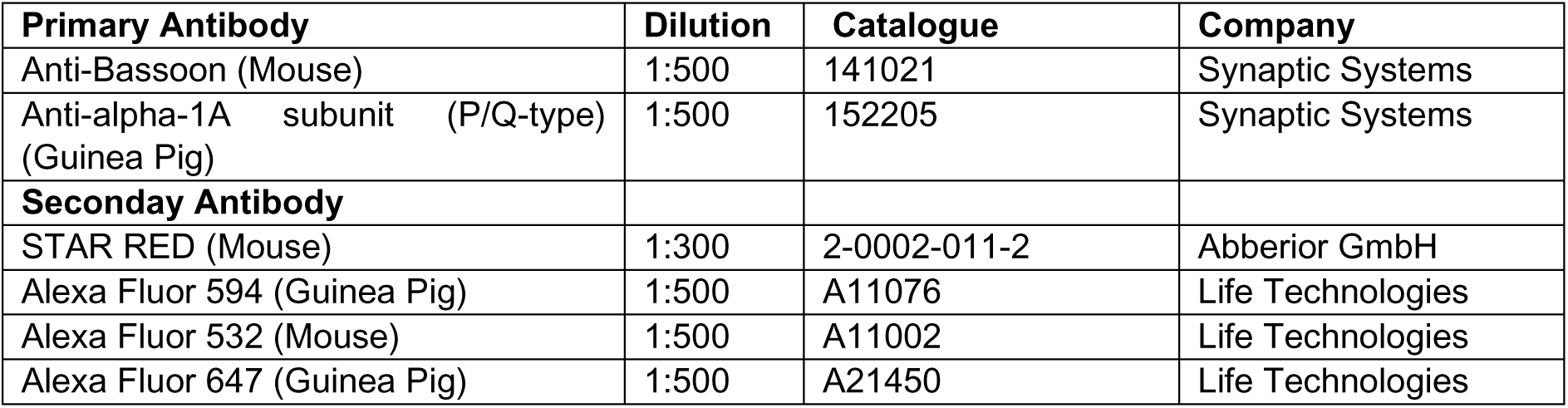

#### Stimulated emission depletion microscopy (STED)

A commercial STED inverted microscope (Abberior Expert Line 775 nm, Abberior Instruments GmbH, Göttingen, Germany) was used to obtain super-resolved images of neurons immunolabelled with the proteins of interest. With a sampling of 15nm, a confocal image and a STED image of the same region were recorded. This system used a pulsed depletion laser at 775 nm and two pulsed excitation lasers at 561 nm and 640 nm ^2,6^

### Single-molecule localization microscopy

18 mm round coverslips from the company (OKO lab, Italy) were used to mount the cells, which were then imaged using direct Stochastic Optical Reconstruction Microscopy (dSTORM) at 37°C in a closed chamber (LudinChamber; Life Imaging Services, Switzerland). The coverslips were mounted on an inverted motorized microscope (IX83 Olympus) equipped with a 100× TIRF, 1.49 NA objective that allowed long acquisition by HILO (highly inclined and laminated optical sheet) illumination. The illumination and acquisition were controlled using MetaMorph software. Fiducial markers were placed using 100 nm Tetraspeck beads (Tetraspeck; Thermo Fisher Scientific, USA).To induce stochastic activation of sparse subsets of molecules in the immunolabeled cells, we used dSTORM buffer, which is an enzymatic oxygen scavenging solution consisting of catalase, TCEP, glycerin, glucose, and glucose oxidase dissolved in Tris-HCl buffer. The optimal density of 0.01–0.04 molecules/μm2 was attained using a 300 mW excitation laser for the photoconversion of Alexa 647 fluorophores from ensemble to single molecule density. Once the optimal density was reached, the illumination laser power was reduced to 150 mW. Five stacks of 4000 frames were obtained at an exposure time of 20 ms, resulting in a total of 20,000 images. For dual color dSTORM, Alexa 532 was used to label Bassoon in combination with Alexa 647 for marking VGCC^1,3,7^..

### Semiautomated detection of functional compartment like presynapse and active zone of an excitatory synapse

The semiautomated detection of functional zones in a synapse was carried out as follows: First, the intensity of the epifluorescence/confocal images of the active zone marker, Bassoon, was thresholded to generate the presynapse mask. Next, a morphometry analysis was performed, and the masks were filtered using various morphological filters, such as length, breadth, and area through the IMA plugin within the MetaMorph software (Molecular Devices) (Kedia et al., 2020a, 2020b; Venkatesan et al., 2020). For the confocal images, fluorescent puncta were thresholded with an area between 0.03 and 3 µm^2^. Similar analysis was performed on super-resolution STED images, where the puncta were thresholded with an area between 0.010 and 1 µm^2^ to generate the mask of the active zone within the presynapse. Finally, parameters like area, average intensity, and total intensity of the presynapse and active zone compartments were computed for both the experimental groups of control and TTX^1,2,4,6^.

### Super-resolution Cluster Analysis

Molecular aggregation clusters (nanodomains) were identified from super-resolved STED images using a custom algorithm written as a plugin supported by MetaMorph (Molecular Devices)^3,8,9^. The Palm Tracer plugin was used to detect nanodomains from STED images. Subsequently, nanodomains were analyzed using a bidimensional Gaussian fitting, from which the principal axis (2.3σ long) and the auxiliary axis (2.3σ short, data not shown) were determined. Fitting was performed on each cluster identified as a domain^1,4,6^. Direct parameters such as the area and intensity of the nanodomain were determined.

### Evaluation of first-order phase transitions within nanoscale condensates

The dSTORM data provided information on the total intensity of the cluster. The intensity of a single molecule was calculated, which allowed the determination of the number of molecules within a single cluster. This was calculated by dividing the total intensity of a cluster by the individual intensity of a single molecule. This served as the raw data, which was used for further analysis. A histogram of the probability distribution of the number of molecules detected inside each cluster was plotted. The probability of occurrence decreased rapidly towards clusters with more molecules, which behaved non-uniformly as supramolecular aggregation patterns. The histograms of the collated cluster n values were normalized to obtain the cluster size distribution in each experimental condition. It was ensured that all had a constant bin size in the Control and TTX dataset for a specific protein (Bassoon/VGCC) and age of the synapse (DIV 8/DIV 15), using the Nyquist criterion for binning. A minimum bin size of 100 was used, and a maximum bin size of 400 was used for different groups to avoid hitting a noise floor.

The data was then fitted for first-order phase transition using the formula for free energy change associated with clustering monomers into aggregates. This formula was previously reported as ΔG=a(n)^2/3^ ± b(n) + c (Narayanan et al., 2019). The term ΔG_surface_=an^2/3^ represents the surface energy of the molecules, and ΔG_bulk_=±bn defines the bulk energy term that depends on the ambient monomer concentration (C_amb_) of Bassoon/VGCC and the saturation concentration (C_sat_) of Bassoon/VGCC at equilibrium with the clustered phase. Since the number of molecules (n) in a domain is a function of morphology and local distribution of molecules within clusters, it relates to the cluster radius (R) in nanometers, as n = (R/1 nm)^3^. The localization events in a cluster scale to the cube of cluster radius by assuming a uniform distribution inside clusters.

ΔG(n) was determined from the distribution of cluster sizes using the equation ΔG = −k_B_T Log(P(n) + Log A [for n < nc]), where k_B_ is the Boltzmann constant and T, the temperature (Kelvin). P(n) represents the relative frequency distribution of the cluster size (n), and nc is the critical cluster size that attains a maximum value of ΔG(nc), which is the nucleation barrier. The balance between negative bulk energy that minimizes a system’s free energy and positive bulk energy that maximizes the system’s free energy leads to a nucleation barrier. Once the nucleation barrier is reached, the cluster formation becomes spontaneous. A positive ΔG (C_amb_ < C_sat_) represents that the cluster is in a sub-saturated state, whereas a negative ΔG (C_amb_ > C_sat_) depicts that the cluster is in a supersaturated state. For a cluster of n molecules, ΔGbulk = Δμn and Δμ = k_B_T Log(C_amb_/C_sat_).

The term “bn” representing the bulk energy ΔGbulk was extracted from the data by subtracting “an2/3” from ΔG. After observing that the resulting curve was linear with a negative slope, indicating a supersaturated system, the inflection points (a, b, c values) obtained from the curve fit of the free energy equation ΔG were used to compute two important parameters: the nucleation barrier (Nc), calculated as (2a/3b)^3^, and the critical cluster radius (Rc), calculated as the cube root of the nucleation barrier (Nc). To ensure consistency across individual cells and the cumulative results, there was a careful comparison of each cell’s a, b, and c values and those cells were selected where these values fell within the interquartile range of the collated cumulative values. This step ensured parity between individual cells and the overall analysis. The methodology for analyzing the first-order phase transition for Bassoon and VGCC was adapted as described previously^7,10^.

### Rank order analysis to determine multiplicative scaling factor

A series of analyses were conducted to determine the multiplicative scaling of average intensity between the control and TTX datasets ^2,5,7^. The data were pooled for the control and TTX conditions from all puncta from all cells to generate rank-ordered plots of TTX versus control fluorescence intensities. From both control and TTX data, an equal number of puncta were randomly chosen, and rank ordering was performed from lowest to highest fluorescence intensity. The TTX data were plotted against the control data, and a linear fit was calculated to estimate the multiplicative scaling factor. To determine the accurate scaling factor, a Python code was generated using Jupyter Notebook 6.0.3 from Anaconda Navigator, following the method used earlier ^11^. The TTX distribution was scaled down by an arbitrary multiplicative factor, and only the scaled values greater than the minimum intensity in the control condition were included, i.e., scaled data = (TTX data/scaling factor). A wide range of hypothetical scaling factors was used, and the K-S test was used to compare the scaled distribution with the control distribution. The scaling factor corresponding to the highest p-value was considered as the most accurate, and this process was repeated 100 times from a sampling value of 100 to obtain the scaling factor.

### Extracted parameters to determine the content of free Bassoon and VGCC molecules along with their spatial localization

A few derived parameters were computed to address the behavior of free and clustered Bassoon/VGCC molecules. The parameters are as follows: individual intensity ratio (IR_i_) (IR_i_ = total intensity of one nanodomain/total intensity of presynapse or active zone) and grouped intensity ratio (IR_g_) (IR_g_ = total intensity of all nanodomains/total intensity of presynapse or active zone). Another parameter called area ratio (area of one nanodomain/area of presynapse or active zone) was used to ascertain the compactness of nanodomains. The spatial spread of these molecules in synapses and AZ was studied by computing the following parameters: Inter-cluster distance (Dist_IC_), which describes the distance between nanodomains; distance from the ROI center (Dist_ROIC_), which describes the distance of a nanodomain from the center of the region of interest (synapse or AZ); and distance between Bassoon and VGCC nanodomains (Dist_BV_) within a synapse. The number of synapses and AZs with nanodomain numbers ranging from 1 to 10 was also computed. Python codes from Anaconda Navigator generated using Jupyter notebook 6.0.3 were used to collate the above data.

### Statistics

GraphPad Prism version 7.04 for Windows, developed by GraphPad Software in La Jolla, California, USA (www.graphpad.com), was utilized for performing statistical analysis. The normality of the data was evaluated using both D’Agostino-Pearson Omnibus and Shapiro-Wilk normality tests. Outliers were removed according to ROUT method in Graphpad Prism. Mean ± SEM was used for presenting normally distributed data, whereas median (IQR 25% to 75% interval) was used for non-normally distributed data. The frequency distributions used for constructing probability density function histograms of cluster sizes and cumulative probability distribution function curves were normalized. The difference between Control and TTX conditions in the free energy ΔG curves was evaluated using the K-S test. Linear regression of bn was performed for both control and TTX conditions to determine the value and slope of the bulk energy component.

### Estimating the peak distribution of the number of molecules in clusters of Bassoon and VGCC using both dSTORM and STED data

The present study utilized a workflow that was previously described to estimate the number of molecules in the synapse, cytomatrix of the active zone (CAZ), and nanodomains within the synapse and CAZ ^1-4^. To generate a frequency distribution plot of the number of Bassoon/VGCC molecules in a nanodomain, the number of Bassoon/VGCC molecules obtained from dSTORM was used. This allowed estimation of the number of molecules present in a typical nanodomain across most synapses by utilizing the highest peak value in the frequency distribution. Next, STED data was employed to estimate the ratios of protein distribution (Bassoon/VGCC) in the synapse, AZ, and nanodomains within the synapse and AZ. To estimate the number of molecules in the synapse, the peak value from the frequency distribution was divided by the individual intensity ratio (IRi) in the synapse from the STED data. To estimate the number of molecules in the AZ, the total intensity of the AZ was divided by the total intensity of the synapse to obtain the ratio of the molecules present in the AZ. This ratio was then multiplied by the estimated number of molecules in the synapse calculated in the previous step to obtain the estimated number of molecules in the AZ.To estimate the number of molecules in the nanodomains of the AZ, the calculated number of molecules in the AZ was multiplied by the STED IRi in the AZ. Once the number of molecules in the whole synapse, AZ, and the nanodomains in synapses and AZ was estimated, the free pools were calculated. The free pool of Bassoon/VGCC in the synapse was obtained by subtracting the number of molecules in the nanodomains from the total number of molecules in the synapse. Similarly, the free pool of Bassoon/VGCC in the AZ was calculated by subtracting the number of molecules in the nanodomains of AZ from the total number of molecules in the AZ.

After estimating the number of molecules present in the nanodomains of synapses and AZ, a histogram of the relative frequency of the number of molecules in a synapse/AZ was created using OriginLab 2023 software. The Multiple Peak fit module under Peaks and Baseline was used to analyze the distribution further. This module identified multiple peaks in the histogram and fitted them with a Gaussian model peak function using the equation y= y0 + (A/(w*sqrt(pi/2)))exp(- 2((x-xc)/w)^2).The estimated peaks with the highest R2 value were selected as the final data. If an additional peak ("n+1" peaks) improved the chi-square by at least 10% from its previous value (fit with "n" peaks) and the changes were in the direction towards the model value of 1, it was added. Otherwise, the addition of another component was not approved, and the model was deemed to fit the data for "n" peaks. The residues of the fit were always used to evaluate the quality of the fit.

### Stochastic simulation of calcium dependent vesicular release at CA3 presynapse

In this study, we utilized a biophysical, stochastic, and spatially explicit model of the CA3 presynapse to simulate calcium dynamics leading to vesicular release in response to an action potential ^12^. The model includes relevant molecules arranged in their physiological concentrations in a simplified cuboidal geometry representing a canonical presynapse with dimensions of 0.5 x 0.5 x 4 µm, as well as a cuboidal ER compartment with dimensions of 0.1 x 0.1 x 3.9 µm. We performed simulations using Monte Carlo cell version 3, which tracks the diffusion of each molecule and carries out the user-specified reactions stochastically. The model incorporates several molecular components, including calcium, voltage-dependent calcium channels (VDCCs), plasma membrane calcium ATPase (PMCA), sarco-endoplasmic reticulum calcium ATPase (SERCA), Ryanodine receptors (RyR), and calbindin-D28k. VDCCs open stochastically in response to the voltage stimulus, causing calcium to rush into the cytosol, where it is buffered by calbindin. RyRs open on binding to calcium ions, releasing calcium out of the ER. The activity of calcium pumps (PMCA, SERCA) maintains the calcium concentration in each compartment^12,13^

- The activity of high-voltage activated P/Q-type VDCCs was modeled using a five-state kinetic scheme described by Bischofberger et al. (2002). The transition rates are voltage-dependent and follow exponential relationships: ai(V) = aioexp(V/Vi) and bi(V) = bioexp(-V/Vi). Additionally, the voltage profile corresponding to an action potential has been described previously^14^.
- The PMCA pumps were modeled as described previously^15^, with the open state transitioning to the closed state through two isomerization steps. The open state exhibits higher ATPase activity and binds to calcium with greater affinity. As a result, calcium is released extracellularly due to weaker binding to the closed state.
- RyR was modeled to have low and high activity states, with calcium ions conducted through the open states in each activity state. The kinetic model used has been decribed previously^16^.
- The four-state model described earlier^17^ was used to model SERCA, but it was implemented in Mcell as a six-state model to incorporate multiple calcium binding steps. The rate constants were adjusted to maintain a steady concentration of 250uM in the ER.
- Calbindin-D28k was modeled to have two binding sites with distinct affinities to the calcium ion, based on the previously validated model^18^.
- Finally, calcium sensors were modeled as described earlier^19^, which incorporates vesicular release through three pathways: calcium-dependent synchronous and asynchronous release, as well as calcium-independent spontaneous release.

The model was set to have a cytosolic calcium concentration of 100 nM and an ER calcium concentration of 250 µM in the absence of activity. The readily releasable pool was considered to have seven vesicles, and VDCCs were positioned within a nanodomain of radius 50 nm for all simulations. For synapses with VGCCs greater or equal to 10, a random placement of channels was generated within the 50 nm cluster. For VGCCs 2-8 and distances 10-100nm, release probability was measured for ten different channel placements, which were generated randomly. The nanodomain were placed at different coupling distances from the readily releasable pool as indicated.

Simulations were performed with a time step of 1 µs, and 1000-5000 trials were done for each configuration, depending on the magnitude of the release probability. Smaller release probabilities required more trials to be estimated accurately. The data obtained from the simulations were resampled, with 1000 samples consisting of 100 trials each. The release probability for each sample was calculated as the proportion of trials with at least one vesicle release within 20 ms after action potential initiation. The mean and standard deviation of the measured values of release probability were then calculated. To study the effect of multiplicative scaling of VGCCs, the number of VGCCs in the nanodomain were scaled by scaling factors of 0.5 to 1.5. The scaling was performed on two sets of synapses, with the nanodomain placed 150 nm and 200 nm away from the release site..For the coupling distance of 150 nm, the number of VGCCs in the original population were between 10 to 60, whereas it ranged from from 10 to 110 for the coupling distance of 200 nm. The range of VGCC numbers for each distance was taken to cover the whole range of release probabilities. Every other factor was kept constant during the scaling.

### Image rendering and language corrections

The schematic representation of imaging and analysis workflows was designed using BioRender under a paid subscription. Freely available OpenAI models were utilized for spelling and grammar corrections in the text.

