## Supplementray Figures for "Nanoscale Lattice Organization of Molecular Condensates Drives Compositional Degeneracy in Synaptic Plasticity"

### Supplementary Figures

Figure S1

Scheme 1

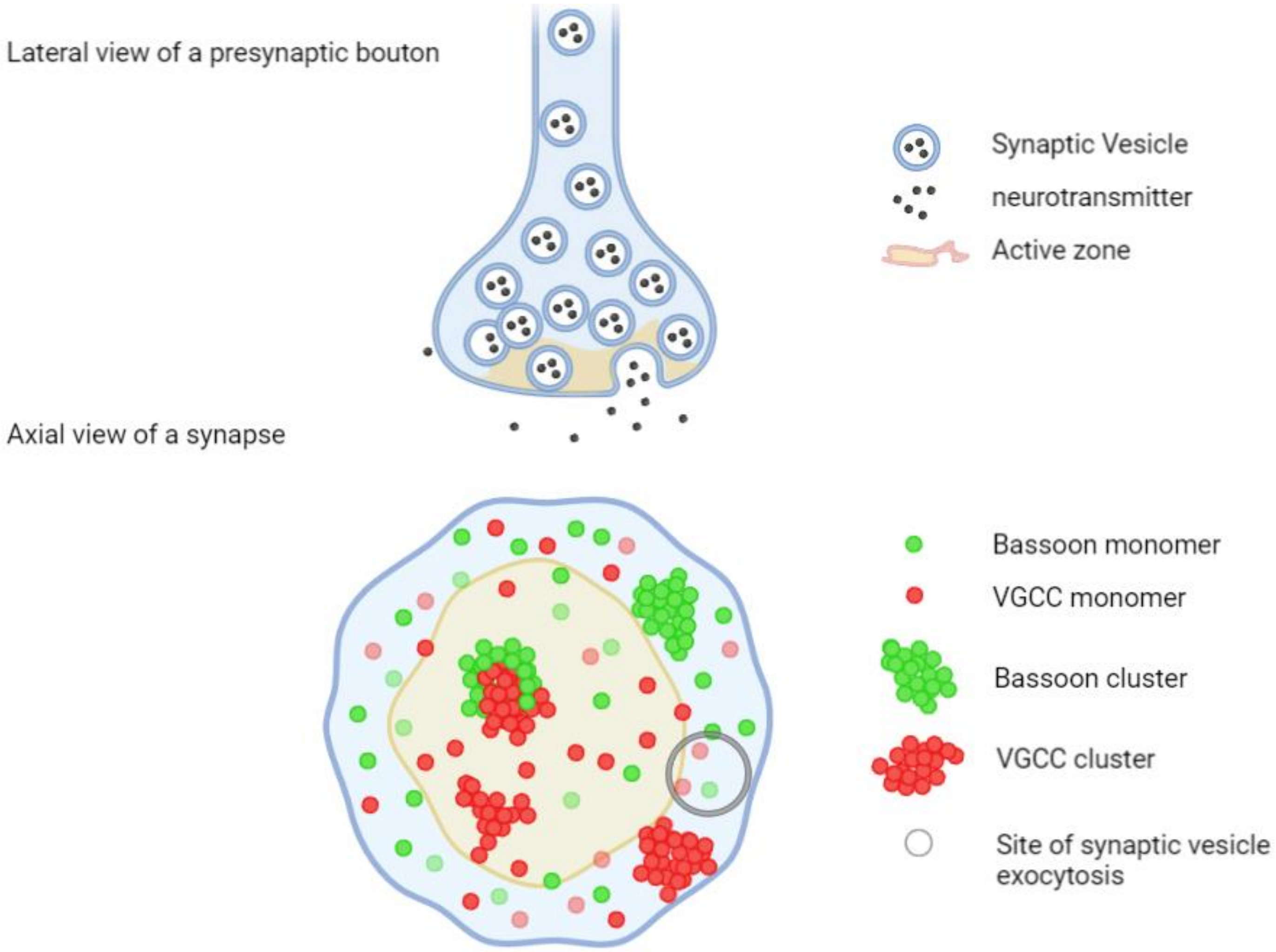


**Figure S1**

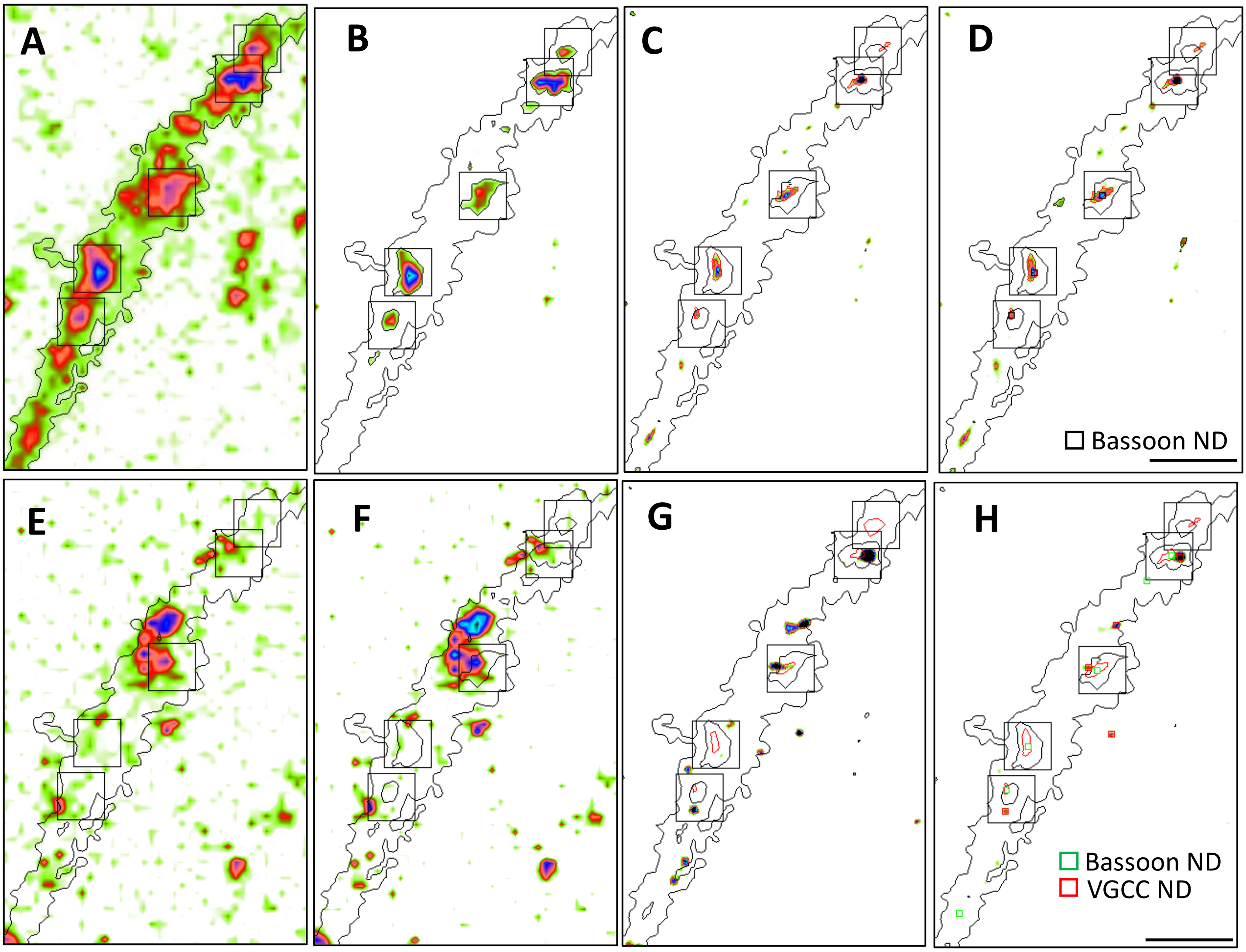


**Figure S2**

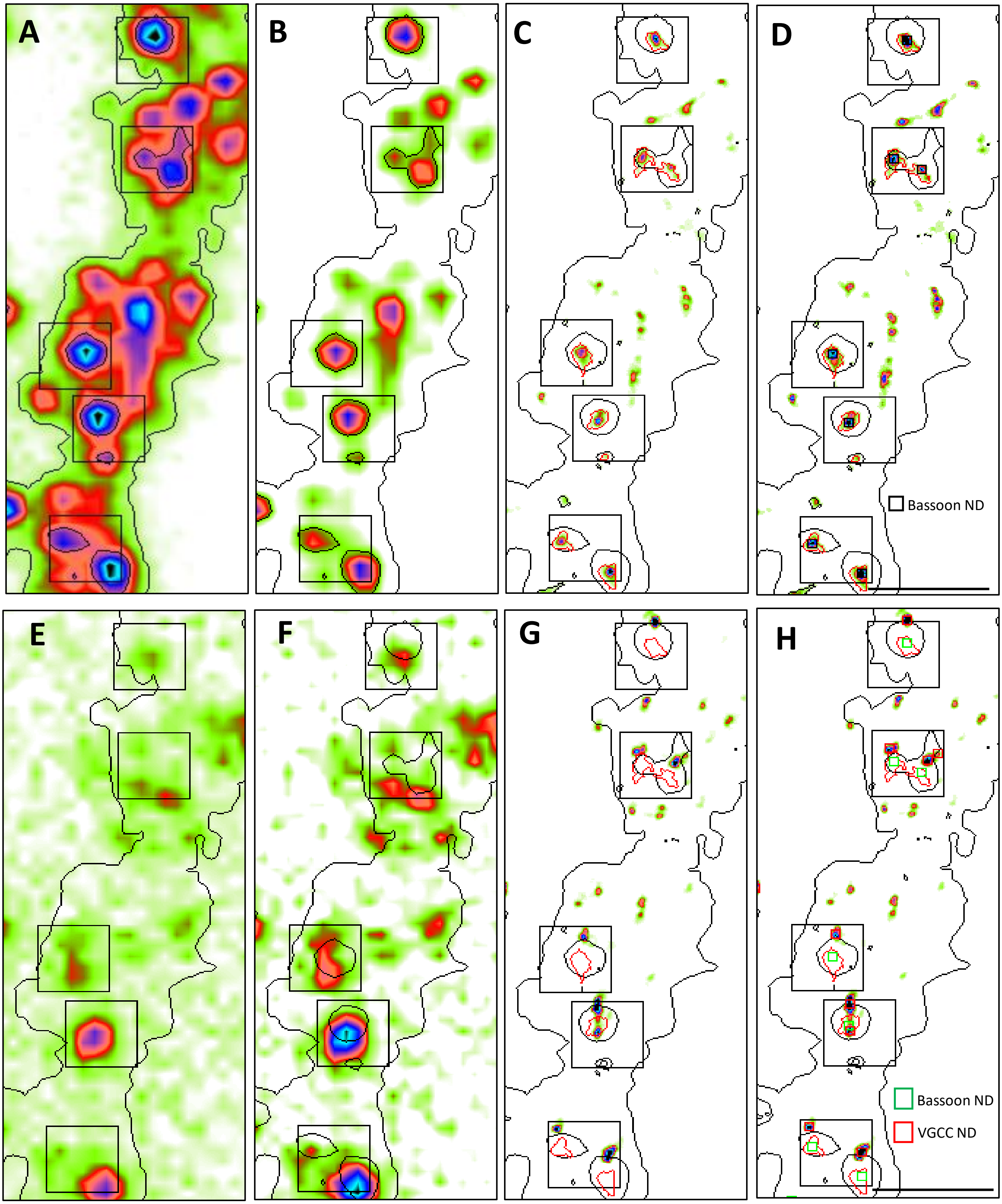


**Figure S3**

Scheme 2

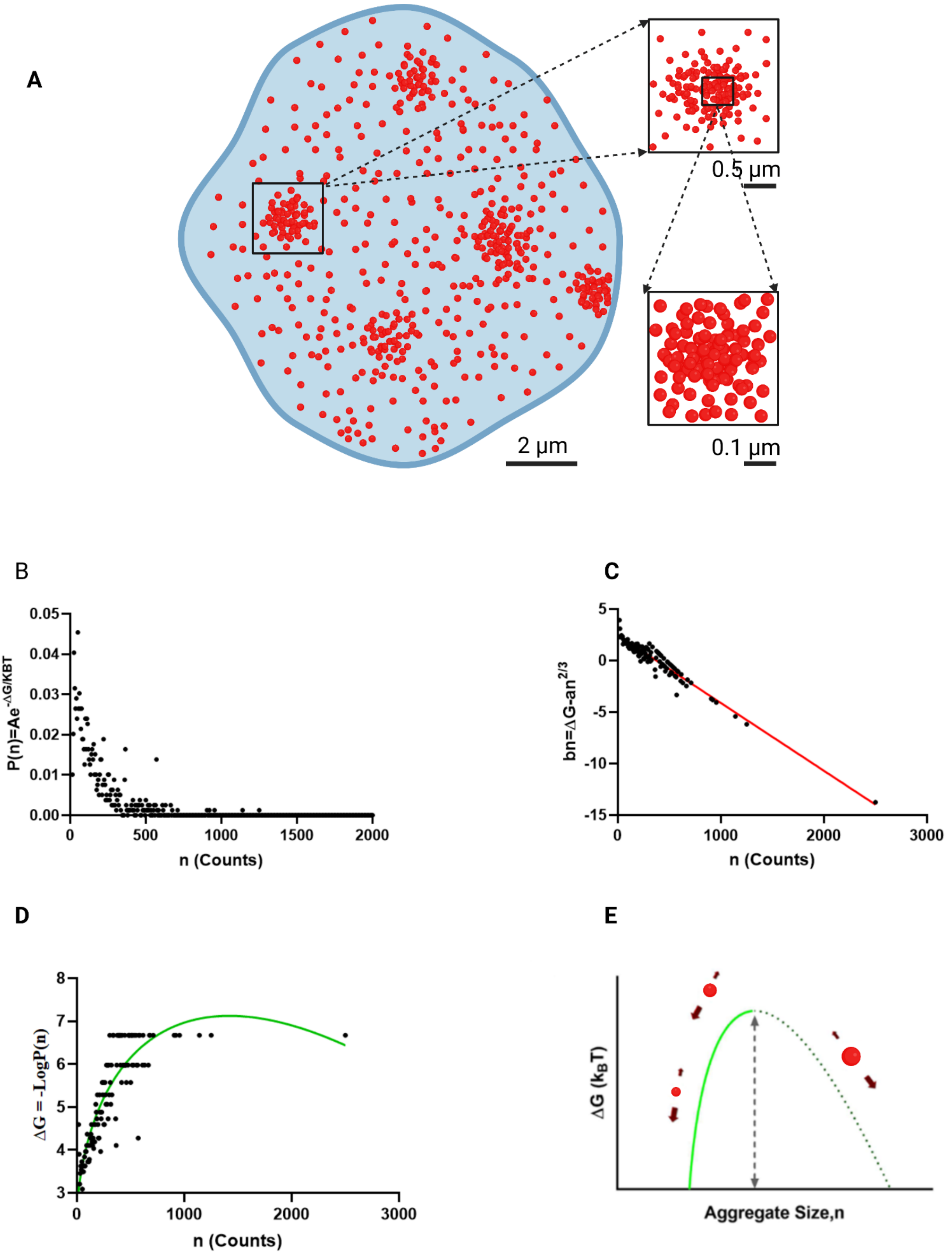


**Figure S4**

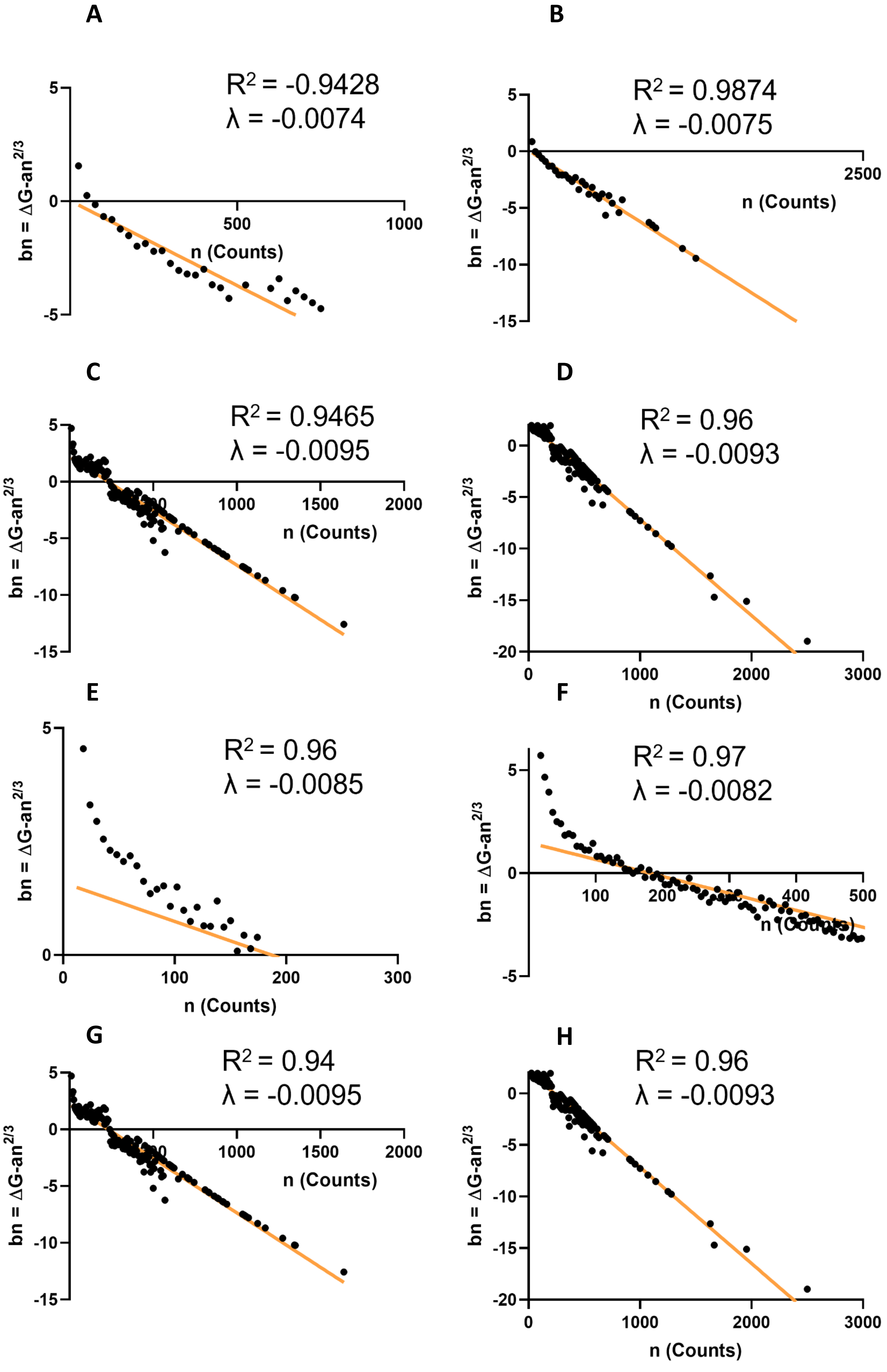

A
R² = -0.9428
λ = -0.0074
bn = ΔG-an^2/3
n (Counts)
B
R² = 0.9874
λ = -0.0075
bn = ΔG-an^2/3
n (Counts)
C
R² = 0.9465
λ = -0.0095
bn = ΔG-an^2/3
n (Counts)
D
R² = 0.96
λ = -0.0093
bn = ΔG-an^2/3
n (Counts)
E
R² = 0.96
λ = -0.0085
bn = ΔG-an^2/3
n (Counts)
F
R² = 0.97
λ = -0.0082
bn = ΔG-an^2/3
n (Counts)
G
R² = 0.94
λ = -0.0095
bn = ΔG-an^2/3
n (Counts)
H
R² = 0.96
λ = -0.0093
bn = ΔG-an^2/3
n (Counts)


**Figure S5**

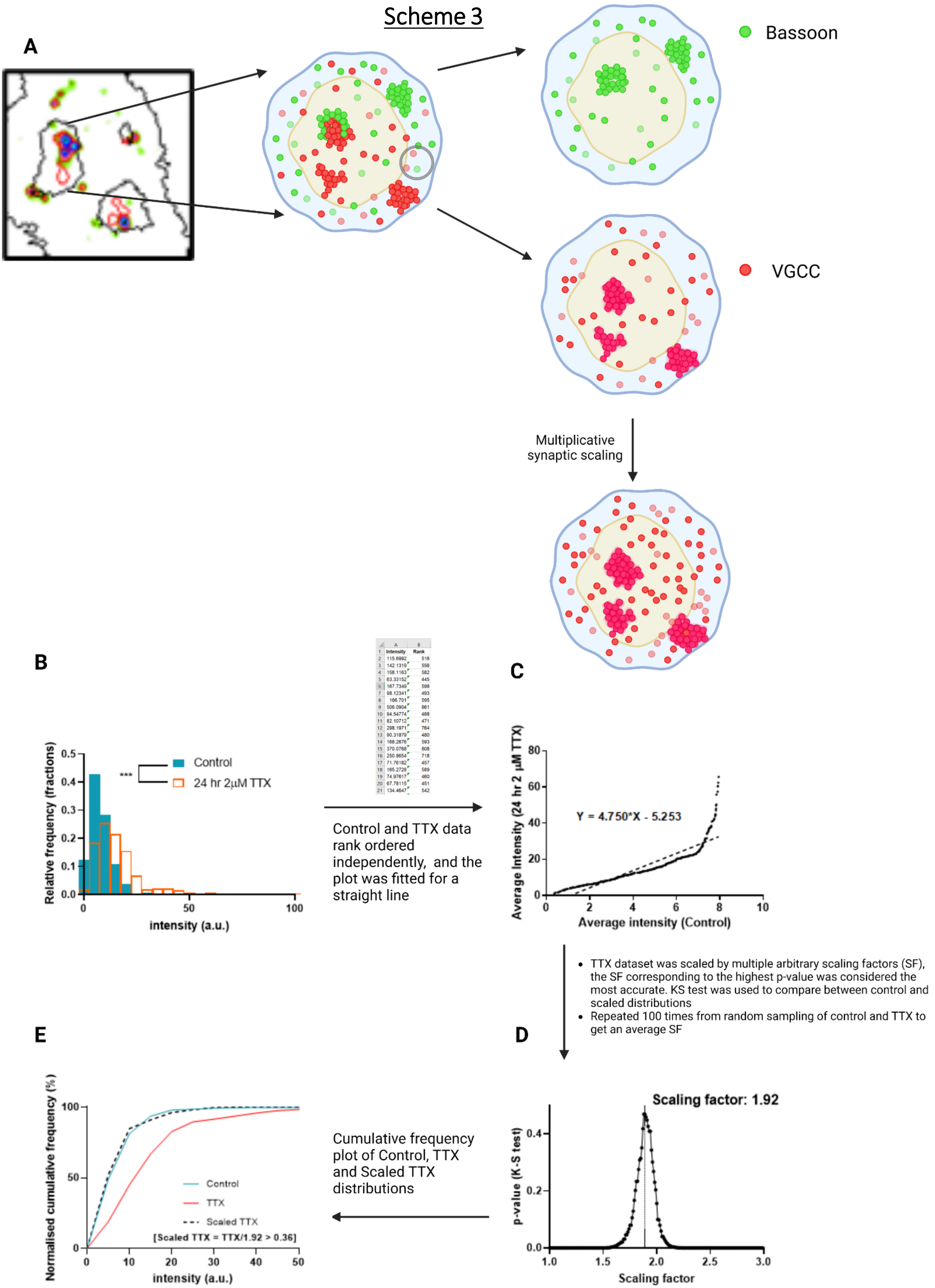
Scheme 3
A
Bassoon
VGCC
Multiplicative synaptic scaling
B
Relative frequency (fractions)
Control
24 hr 2μM TTX
***
intensity (a.u.)
Control and TTX data rank ordered independently, and the plot was fitted for a straight line
C
Average Intensity (24 hr 2 μM TTX)
Y = 4.750*X - 5.253
Average intensity (Control)
• TTX dataset was scaled by multiple arbitrary scaling factors (SF), the SF corresponding to the highest p-value was considered the most accurate. KS test was used to compare between control and scaled distributions
• Repeated 100 times from random sampling of control and TTX to get an average SF
D
Scaling factor: 1.92
p-value (K-S test)
Scaling factor
E
Normalised cumulative frequency (%)
Control
TTX
Scaled TTX
[Scaled TTX = TTX/1.92 > 0.36]
intensity (a.u.)
Cumulative frequency plot of Control, TTX and Scaled TTX distributions

### Figure S6

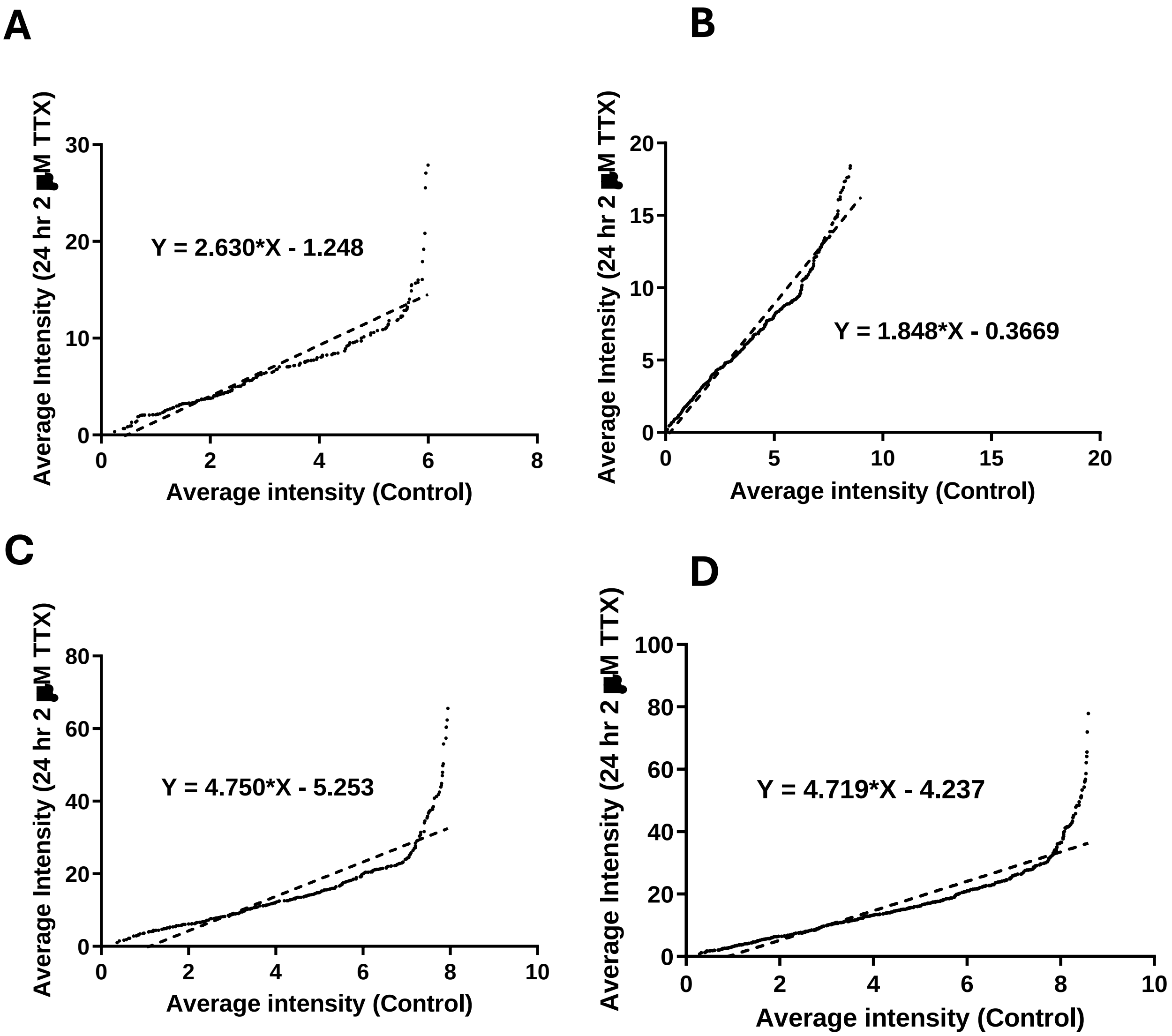
A
Y = 2.630*X - 1.248
Average Intensity (24 hr 2 ■M TTX)
Average intensity (Control)
B
Y = 1.848*X - 0.3669
Average Intensity (24 hr 2 ■M TTX)
Average intensity (Control)
C
Y = 4.750*X - 5.253
Average Intensity (24 hr 2 ■M TTX)
Average intensity (Control)
D
Y = 4.719*X - 4.237
Average Intensity (24 hr 2 ■M TTX)
Average intensity (Control)

**Figure S7**

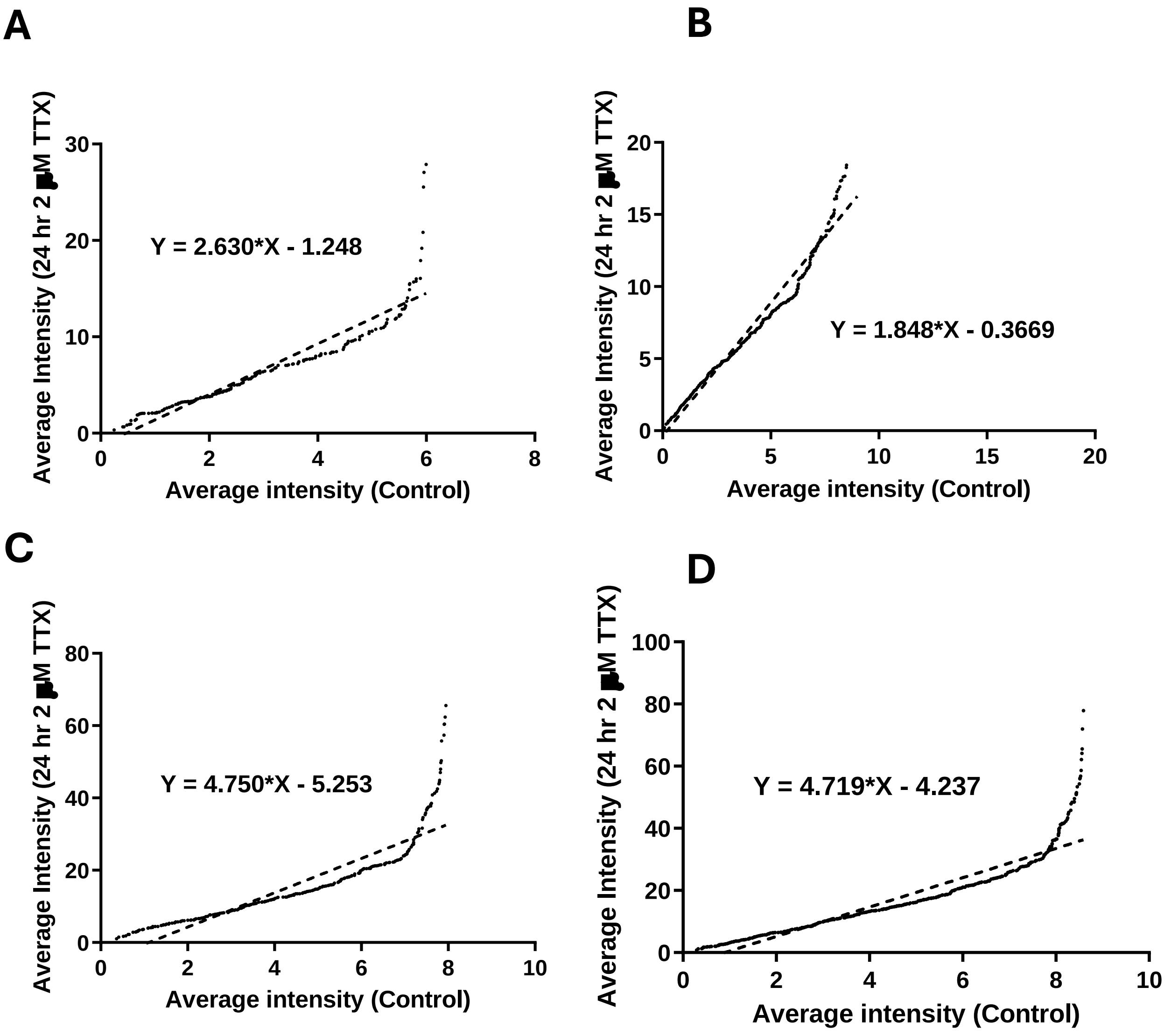

A
Y = 2.630*X - 1.248
Average intensity (Control)
B
Y = 1.848*X - 0.3669
Average intensity (Control)
C
Y = 4.750*X - 5.253
Average intensity (Control)
D
Y = 4.719*X - 4.237
Average intensity (Control)

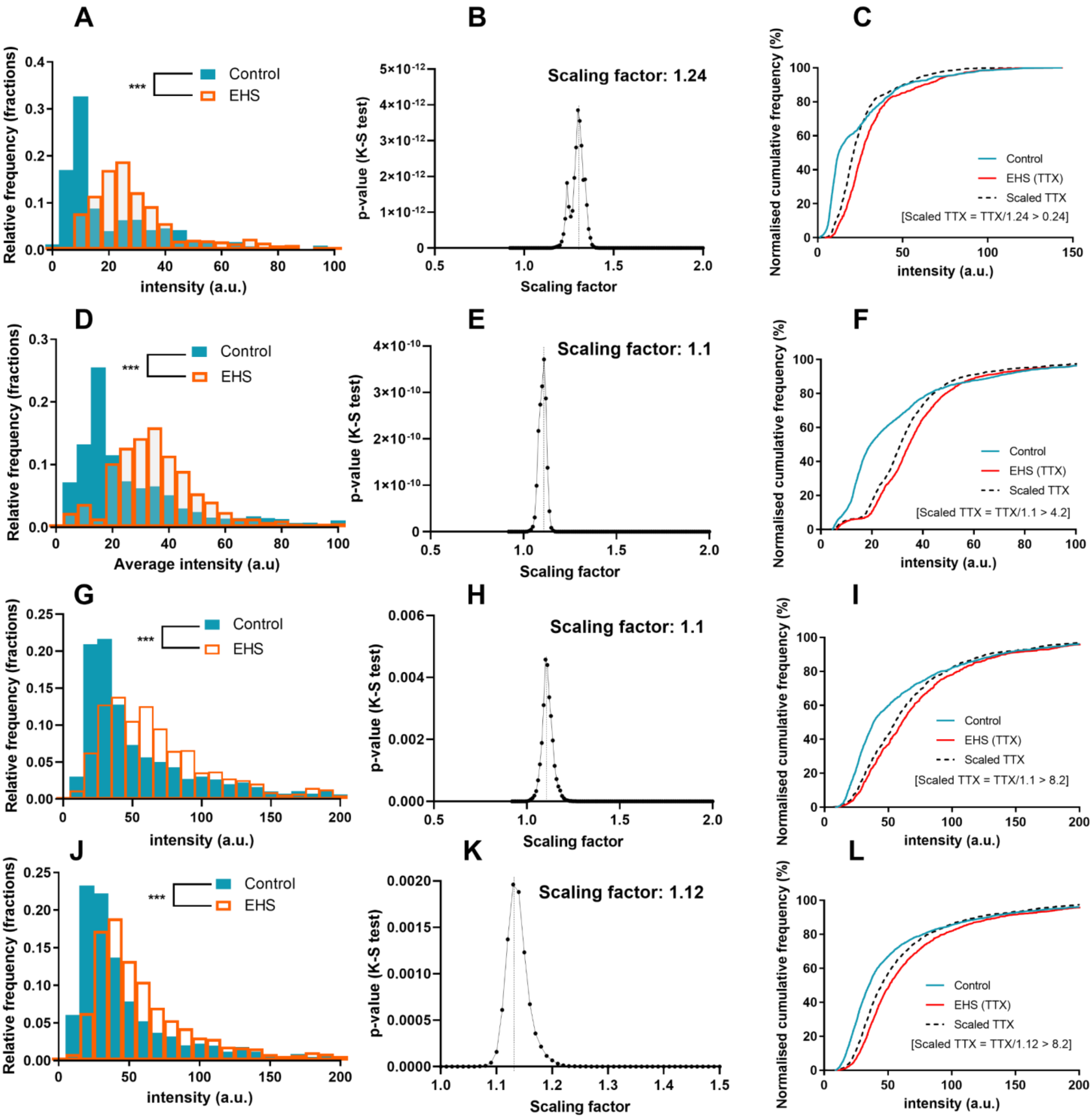
A
B
C
D
E
F
G
H
I
J
K
L
Control
EHS
***
Relative frequency (fractions)
intensity (a.u.)
Average intensity (a.u)
p-value (K-S test)
Scaling factor
Scaling factor: 1.24
Scaling factor: 1.1
Scaling factor: 1.12
Normalised cumulative frequency (%)
EHS (TTX)
Scaled TTX
[Scaled TTX = TTX/1.24 > 0.24]
[Scaled TTX = TTX/1.1 > 4.2]
[Scaled TTX = TTX/1.1 > 8.2]
[Scaled TTX = TTX/1.12 > 8.2]

**Figure S8**

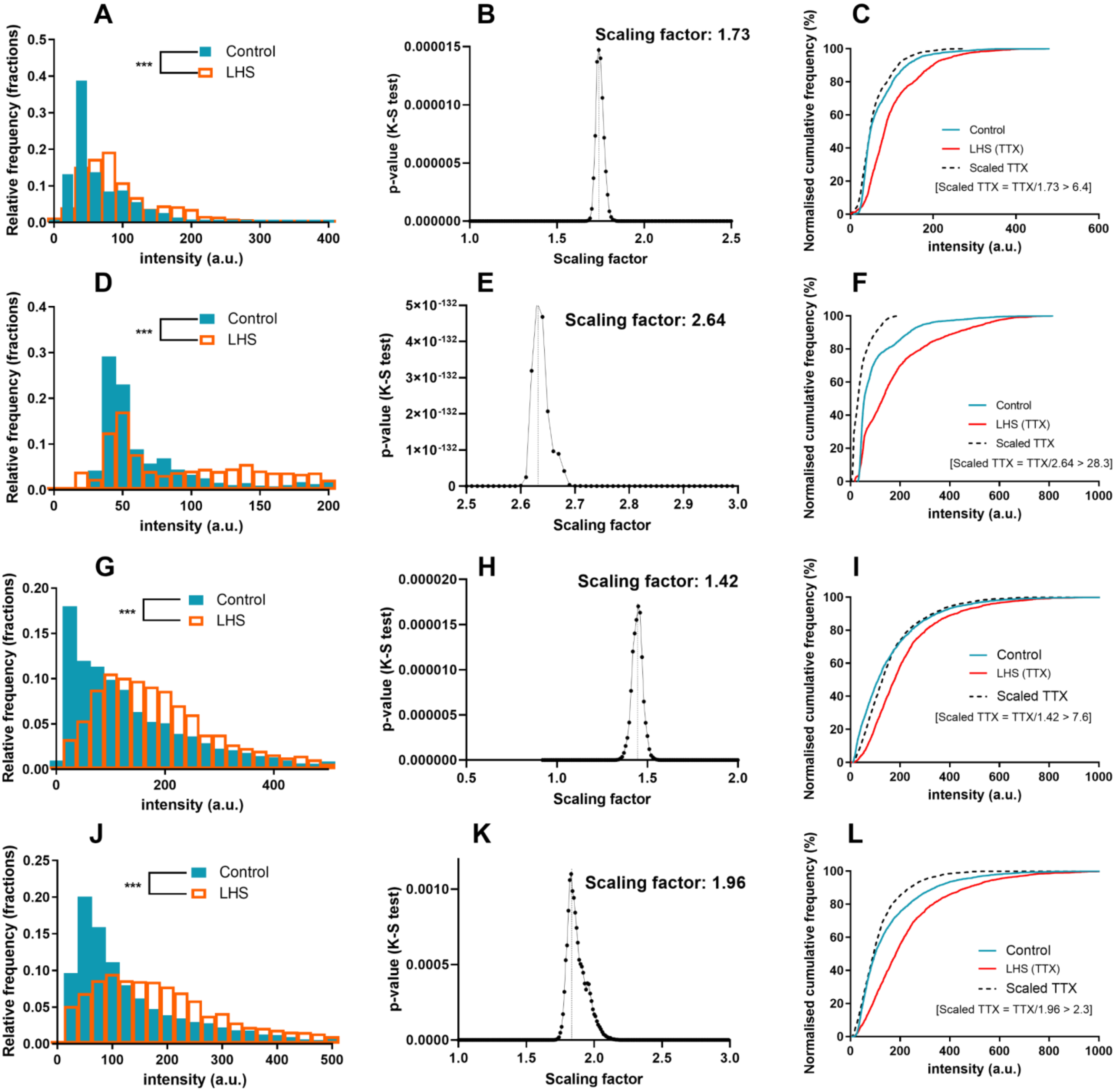
A
Control
LHS
***
Relative frequency (fractions)
intensity (a.u.)
B
Scaling factor: 1.73
p-value (K-S test)
Scaling factor
C
Control
LHS (TTX)
Scaled TTX
[Scaled TTX = TTX/1.73 > 6.4]
Normalised cumulative frequency (%)
intensity (a.u.)
D
E
Scaling factor: 2.64
F
[Scaled TTX = TTX/2.64 > 28.3]
G
H
Scaling factor: 1.42
I
[Scaled TTX = TTX/1.42 > 7.6]
J
K
Scaling factor: 1.96
L
[Scaled TTX = TTX/1.96 > 2.3]

### Figure S9

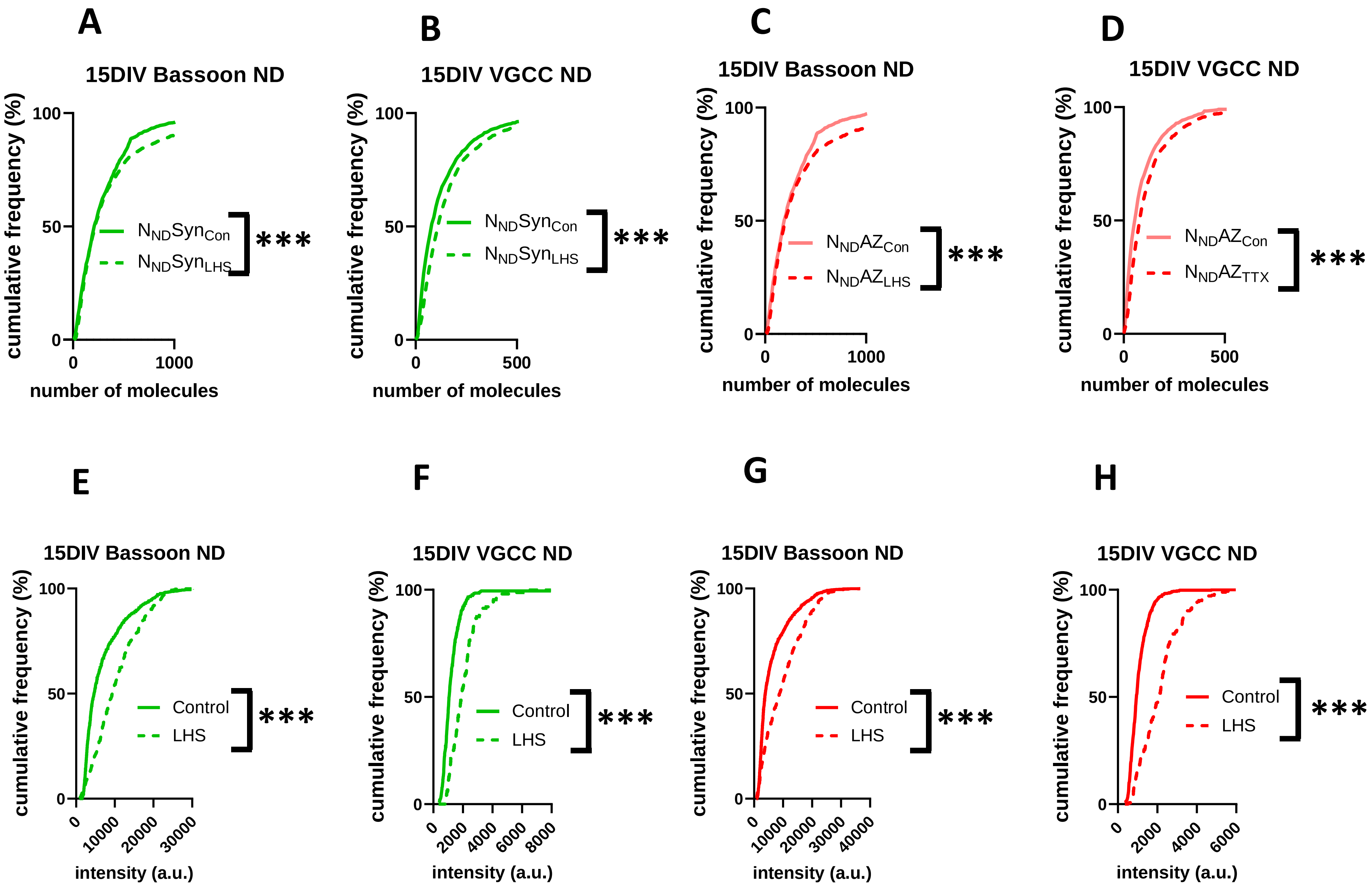

A
15DIV Bassoon ND
cumulative frequency (%)
number of molecules
$N_{ND}Syn_{Con}$
$N_{ND}Syn_{LHS}$
***
B
15DIV VGCC ND
cumulative frequency (%)
number of molecules
$N_{ND}Syn_{Con}$
$N_{ND}Syn_{LHS}$
***
C
15DIV Bassoon ND
cumulative frequency (%)
number of molecules
$N_{ND}AZ_{Con}$
$N_{ND}AZ_{LHS}$
***
D
15DIV VGCC ND
cumulative frequency (%)
number of molecules
$N_{ND}AZ_{Con}$
$N_{ND}AZ_{TTX}$
***
E
15DIV Bassoon ND
cumulative frequency (%)
intensity (a.u.)
Control
LHS
***
F
15DIV VGCC ND
cumulative frequency (%)
intensity (a.u.)
Control
LHS
***
G
15DIV Bassoon ND
cumulative frequency (%)
intensity (a.u.)
Control
LHS
***
H
15DIV VGCC ND
cumulative frequency (%)
intensity (a.u.)
Control
LHS
***


Figure S10

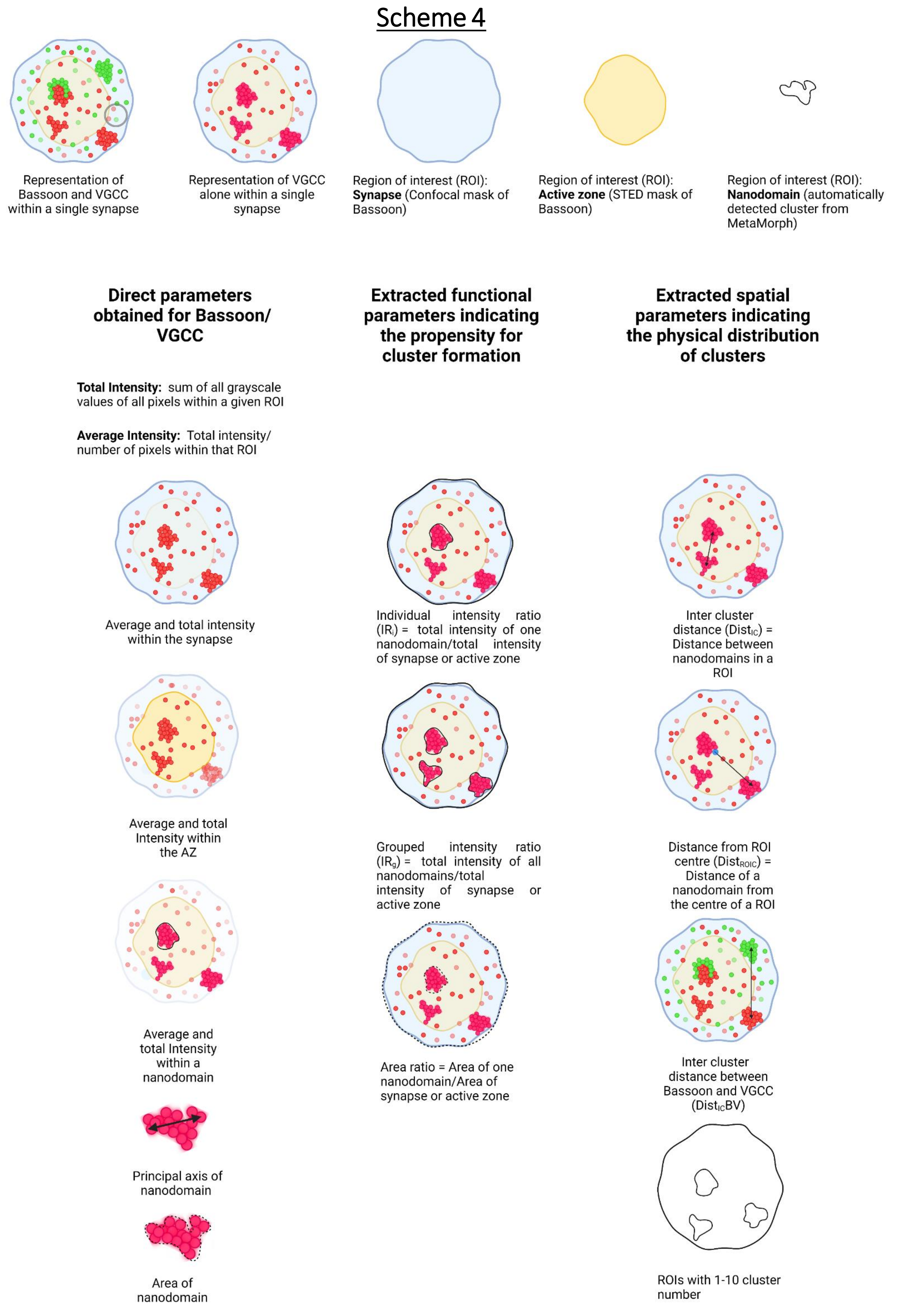
Scheme 4
Representation of Bassoon and VGCC within a single synapse
Representation of VGCC alone within a single synapse
Region of interest (ROI): **Synapse** (Confocal mask of Bassoon)
Region of interest (ROI): **Active zone** (STED mask of Bassoon)
Region of interest (ROI): **Nanodomain** (automatically detected cluster from MetaMorph)
**Direct parameters obtained for Bassoon/ VGCC**
**Extracted functional parameters indicating the propensity for cluster formation**
**Extracted spatial parameters indicating the physical distribution of clusters**
**Total Intensity:** sum of all grayscale values of all pixels within a given ROI
**Average Intensity:** Total intensity/ number of pixels within that ROI
Average and total intensity within the synapse
Individual intensity ratio ($IR_i$) = total intensity of one nanodomain/total intensity of synapse or active zone
Inter cluster distance ($Dist_{IC}$) = Distance between nanodomains in a ROI
Average and total Intensity within the AZ
Grouped intensity ratio ($IR_g$) = total intensity of all nanodomains/total intensity of synapse or active zone
Distance from ROI centre ($Dist_{ROIC}$) = Distance of a nanodomain from the centre of a ROI
Average and total Intensity within a nanodomain
Area ratio = Area of one nanodomain/Area of synapse or active zone
Inter cluster distance between Bassoon and VGCC ($Dist_{IC}BV$)
Principal axis of nanodomain
Area of nanodomain
ROIs with 1-10 cluster number

**Figure S11**

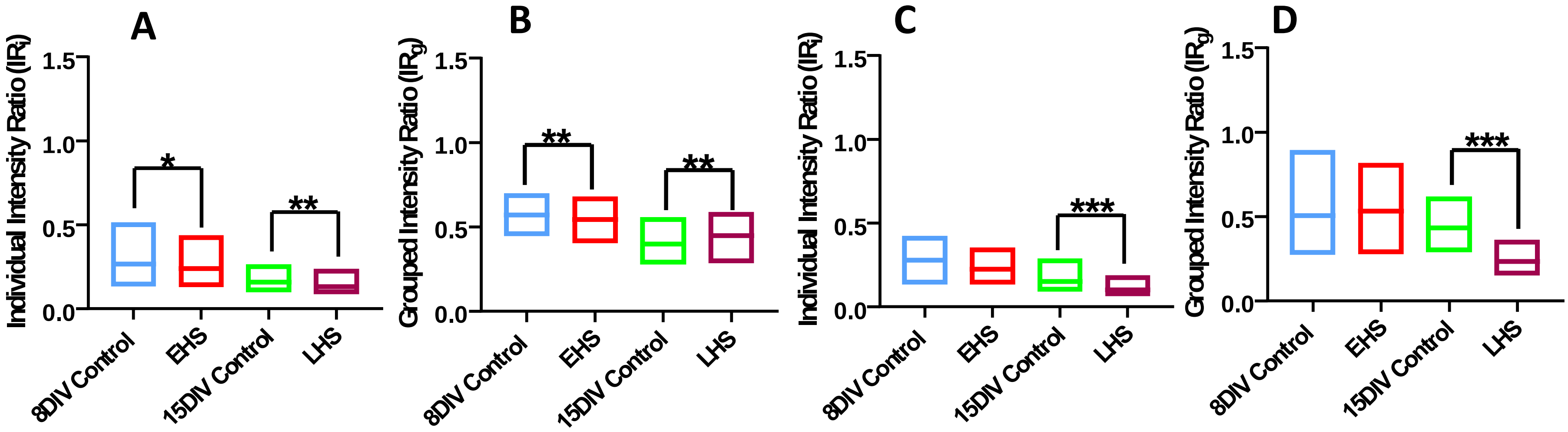
A
B
C
D
Individual Intensity Ratio (IRi)
Grouped Intensity Ratio (IRg)
1.5
1.0
0.5
0.0
8DIV Control
EHS
15DIV Control
LHS
*
**
***

Figure S12

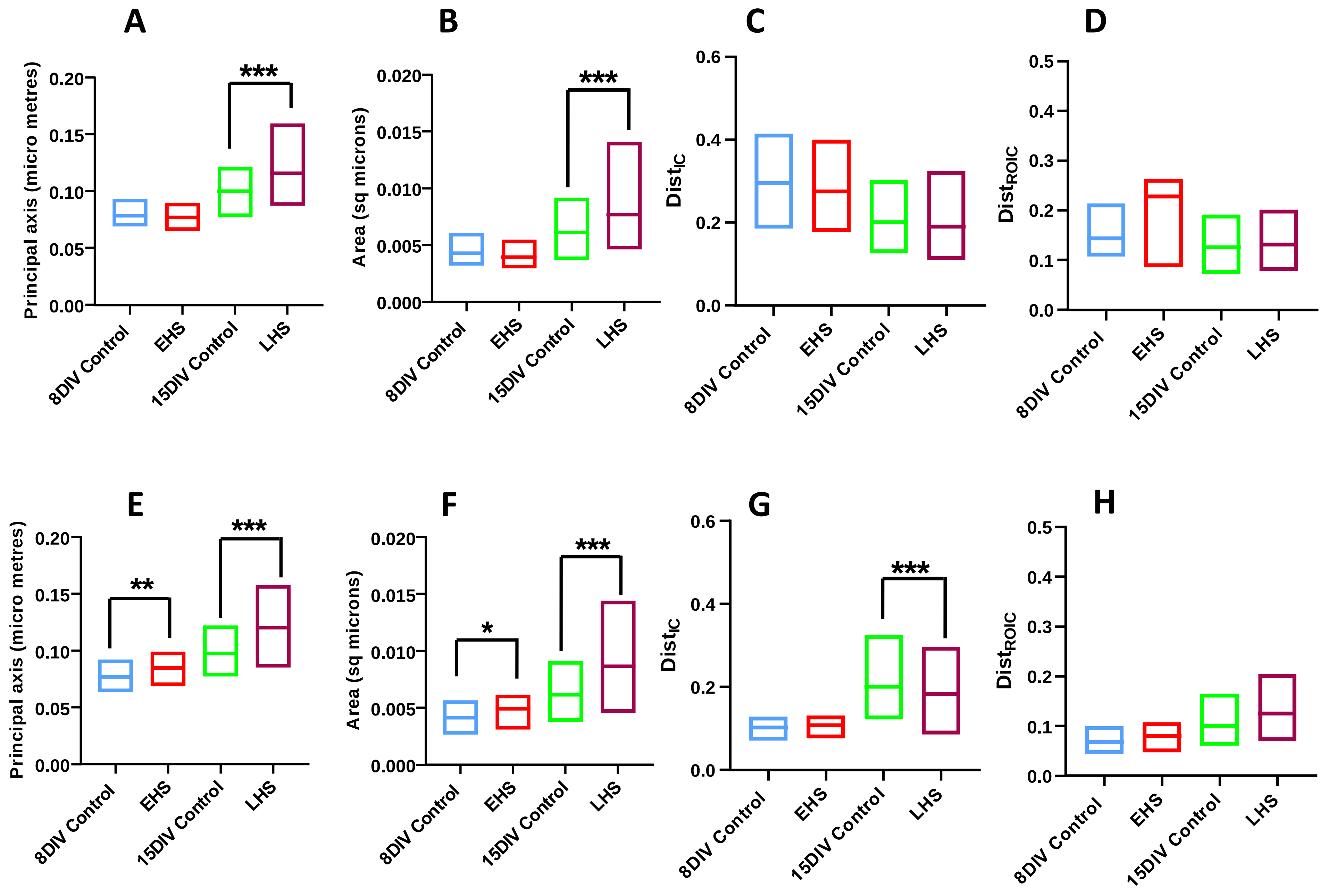
A
B
C
D
E
F
G
H
Principal axis (micro metres)
Area (sq microns)
$Dist_{IC}$
$Dist_{ROIC}$
8DIV Control
EHS
15DIV Control
LHS
***
**
*

Figure S13

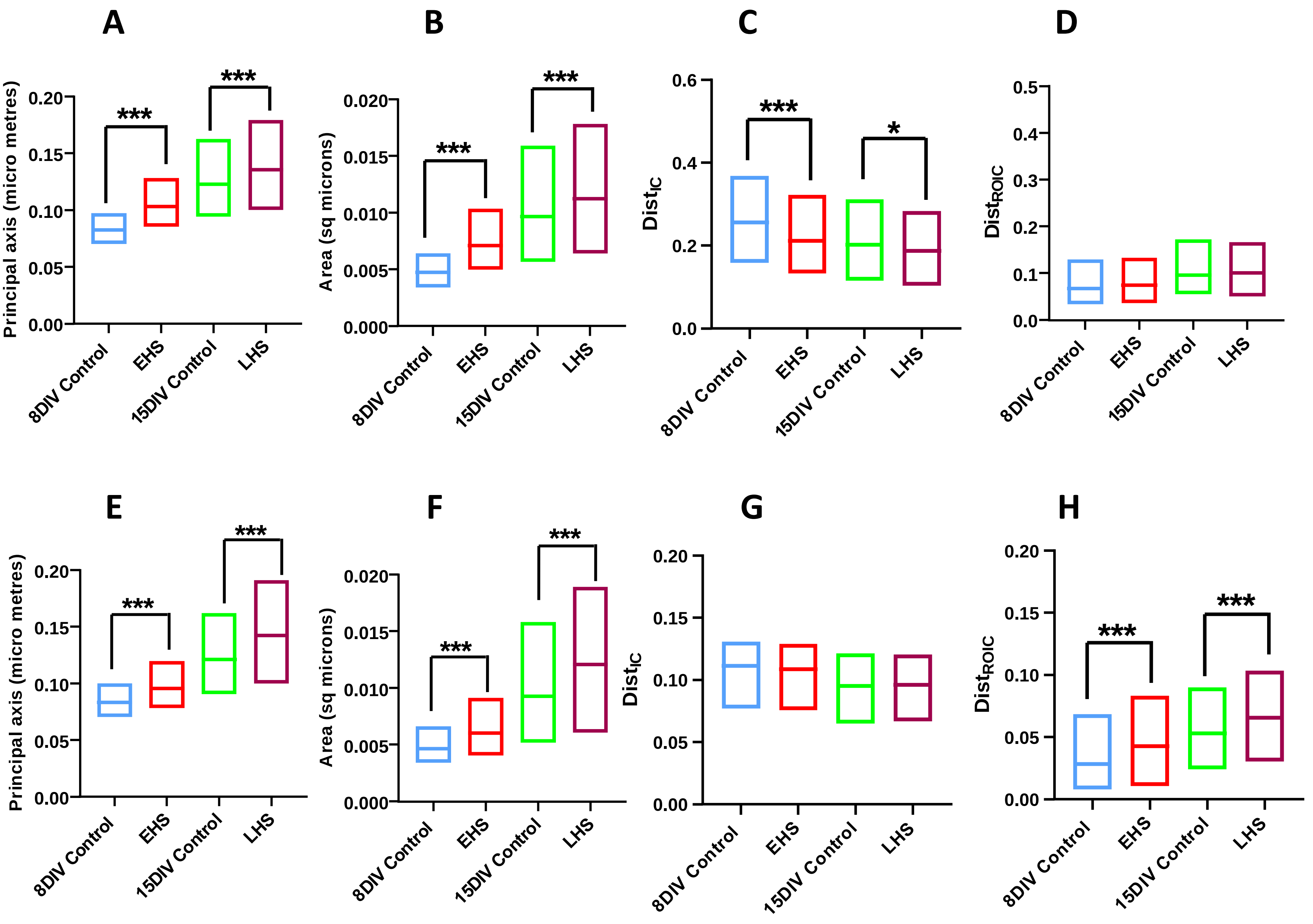
A
B
C
D
E
F
G
H
Principal axis (micro metres)
Area (sq microns)
Dist$_{IC}$
Dist$_{ROIC}$
8DIV Control
EHS
15DIV Control
LHS
***
*

Figure S14
