## Supplementary Table for "Nanoscale Lattice Organization of Molecular Condensates Drives Compositional Degeneracy in Synaptic Plasticity"

| | **8DIV Scaling factors** | | | | **15DIV Scaling factors** | | | |
|---|---|---|---|---|---|---|---|---|
| | Synapse | AZ | Synapse ND | AZ ND | Synapse | AZ | Synapse ND | AZ ND |
| **Bassoon** | 1.24*** | 1.11*** | 1.1*** | 1.12*** | 1.73*** | 2.64*** | 1.42*** | 1.96*** |
| **VGCC** | 1.62 | 1.47 | 1 | 1.15 | 1.92 | 1.92 | 1.37 | 1.41 |

| | Control (λ) | TTX (λ) |
|---|---|---|
| 8DIV Bassoon | -0.0032 | -0.0065 |
| 8DIV VGCC | -0.0011 | -0.0038 |
| 15DIV Bassoon | -0.0117 | -0.0094 |
| 15DIV VGCC | -0.0054 | -0.0067 |
